# Multimodal allostery in a single-domain protein

**DOI:** 10.64898/2026.09.14.750605

**Authors:** Yujing Ma, Qile Jin, Ben Lehner, Chenchun Weng

**Affiliations:** Department of Pathology, The First Affiliated Hospital of USTC, Division of Life Sciences and Medicine, University of Science and Technology of China, Hefei, Anhui, 230036, China; Intelligent Pathology Institute, Division of Life Sciences and Medicine, University of Science and Technology of China, Hefei, Anhui, 230036, China; Wellcome Sanger Institute, Wellcome Genome Campus, Hinxton, United Kingdom; Centre for Genomic Regulation (CRG), The Barcelona Institute of Science and Technology, Barcelona, Spain; Universitat Pompeu Fabra (UPF), Barcelona, Spain; Institució Catalana de Recerca i Estudis Avançats (ICREA), Barcelona, Spain

## Abstract

Allosteric communication between distant sites in proteins is central to biological regulation and underlies the efficacy of many drugs. Allostery allows proteins to function as molecular switches, with their activities controlled by ligand binding, covalent modifications, and mutations outside their active sites. Allosteric maps have been constructed for several proteins, but these all quantify regulation of a single site in each protein. How the allosteric maps of different sites compare is unclear. Here, we address this question by charting complete allosteric maps for two structurally distant binding sites in the oncoprotein KRAS. Unexpectedly, these maps reveal three different forms of allosteric control, that we term coupled, anti-coupled and independent allostery. Allostery in this small protein is therefore complex and multimodal. The co-existence of multiple allosteric networks in a single-domain protein has important conceptual implications and, if widespread, may allow the development of new classes of therapeutics that not only modulate targets but also tune their functional outputs.

## Introduction

Energetic coupling between distant sites in proteins is central to biological control, allowing proteins to be regulated by perturbations outside of their active sites(*1–3*). Allostery allows regulation by post-translational modifications, small molecule binding, and interactions with other macromolecules(*4–6*). Allostery is also an important cause of genetic disease, for example underlying the pathogenicity of many activating driver mutations in oncogenes(*5*, *7*) and loss-of-function germline variants (*5*, *8–11*). Conversely, many highly-effective drugs have allosteric mechanisms, binding outside of protein active sites to inhibit or modulate activity(*5*, *12–17*).

Allostery is widespread and may be an intrinsic property of all proteins(*3*, *4*, *11*, *18–21*). Recently, using mutations, it has become possible to construct complete maps of allosteric coupling to protein active sites(*4*, *11*, *19*, *22–25*). These comprehensive maps have revealed that allostery is pervasive, distance-dependent, and variable in strength across different protein regions. Allosteric mapping has identified novel allosteric sites on protein surfaces, including potentially druggable surface pockets(*23*, *25*). Comparing allosteric coupling in homologous proteins has shown that allostery evolves quite quickly, with each protein in a family having a unique allosteric surface to therapeutically target(*24*, *26*).

To date, allosteric maps have only been charted for a single active site in each protein. Many proteins are, however, multi-functional. For example, many human proteins bind multiple interaction partners through structurally distinct interfaces(*27*). Allosteric regulation of two different binding sites in a protein could be coupled (i.e. positively or negatively correlated) or independent. Positively correlated allostery could occur if a protein has two different conformational or energetic states and two binding partners bind the same state. In contrast, anti-correlated allostery could occur if two proteins bind mutually exclusive states. Independent allostery could result if two proteins bind energetically-independent states or are regulated by distinct allosteric networks.

The onco-protein KRAS provides an attractive model for studying allosteric regulation(*28*, *29*). KRAS is one of the most frequently mutated genes in human cancers and a classic allosteric switch(*7*). KRAS binds diverse interaction partners through overlapping and distinct interfaces, with binding through the main effector interface promoted or inhibited depending upon whether the protein is bound to GTP or GDP(*30–35*). We previously constructed six complete allosteric maps for KRAS, quantifying how mutations throughout the protein change binding to six different proteins that bind to the effector interface(*23*). The six allosteric maps are highly similar, with most mutations having correlated effects on all six interactions. A small number of mutations do, however, have more specific effects, differentially affecting binding to one or a subset of the six proteins. This was most evident for a GDP-state-favoured interaction partner, where a set of mutations had anti-correlated effects relative to binding to the other five proteins(*23*).

Here we use KRAS as a model to comprehensively quantify allosteric regulation of two structurally-distinct interfaces in a protein. Quantifying thousands of binding free energy changes for mutations reveals that three different modes of allosteric regulation co-exist in one protein that we term coupled, anti-coupled and independent allostery. Allostery in in a small single-domain protein is therefore complex, with multiple overlapping allosteric networks.

## Results

### Energetic and allosteric maps for two KRAS interfaces

To globally compare allosteric regulation of two structurally distinct binding interfaces in the same protein domain we quantified the energetic effects of all amino acid (aa) substitutions on the binding of KRAS to proteins through two interfaces (Fig. 1A–D). The first interface is the effector interface, located in the switch-I region of KRAS (approximately residues I24–R41)(*36*), which binds the RAS-binding domain (RBD) of RAF1 and other effector proteins(*31*, *32*, *36*, *37*), as well as laboratory selected affinity reagents, including the Designed Ankyrin Repeat Proteins (DARPins) K27 and K55(*35*). We refer to this interface as ‘interface 1’. The second interface (‘interface 2’) is located on the α3/loop 7/α4 region of KRAS and is bound by two additional DARPins, K13 and K19(*34*). This surface partially overlaps the proposed α3–α4 dimerization interface and is also bound by additional engineered binders, including the monobody 12D1/12D5 and affimer K3(*34*, *38*) (Fig. 1D). Most proteins binding the effector interface have higher affinity for GTP-bound KRAS but DARPin K27 binds with higher affinity to GDP-KRAS(*34*). K13 and K19 bind interface 2 with similar affinity for GTP- and GDP-bound KRAS(*34*), already suggesting the existence of independent allosteric regulation of interface 1.

**Figure 1.**
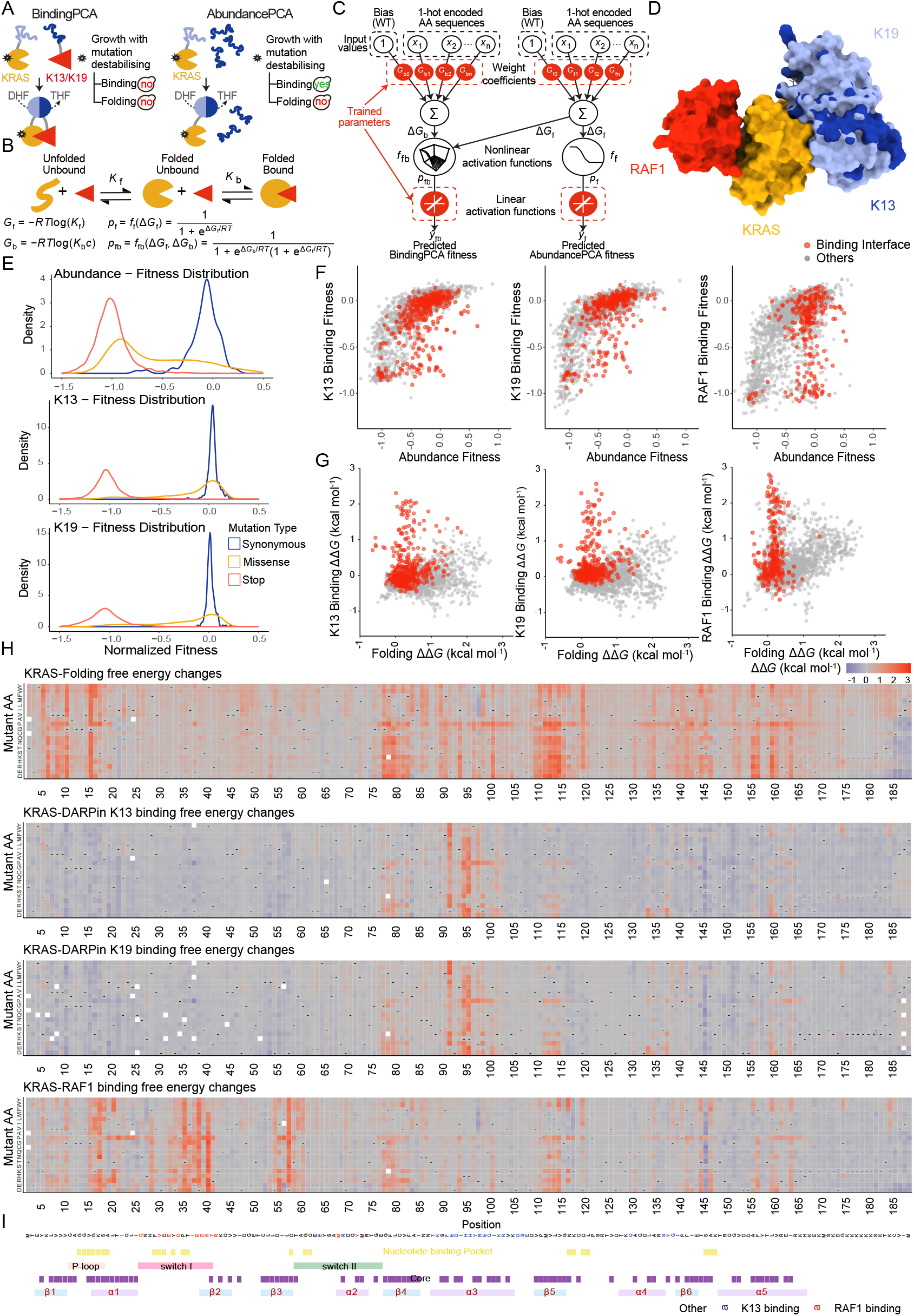
Charting complete energetic and allosteric maps for two binding interfaces of KRAS. (A) Overview of the ddPCA screening strategy. “Yes” indicates normal yeast growth; “No” indicates defective growth. DHF, dihydrofolate; THF, tetrahydrofolate. (B) Three-state equilibrium and corresponding thermodynamic model. Δ*G*_f_, folding Gibbs free energy; Δ*G*_b_, binding Gibbs free energy; *K*_f_, folding equilibrium constant; *K*_b_, binding equilibrium constant; *c*, concentration of binding partner; *p*_f_, fraction folded; *p*_fb_, fraction folded and bound; *f*_f_, nonlinear function of Δ*G*_f_; *f*_fb_, nonlinear function of Δ*G*_f_ and Δ*G*_b_; *R*, gas constant; *T*, temperature in Kelvin. (C) Neural network architecture used to fit the thermodynamic model to ddPCA data (bottom: target versus output; top: input values) to infer causal changes in folding and binding free energies for single amino acid substitutions. AA, amino acid; WT, wild type. (D) KRAS 3D structures in complex with RAF1-RBD, DARPin K13, and DARPin K19 (PDB ID: 6VJJ, 6H46, and 6H47). (E) Normalized fitness distributions of different mutation types across three library blocks (KRAS-Abundance, KRAS-K13, KRAS-K19). Colors indicate mutation type: blue, synonymous; orange, missense; red, nonsense. (F) Scatter plots comparing abundance and binding fitness for single amino acid substitutions. Interface residues are highlighted in red. (G) Scatter plots comparing binding versus folding free energy changes for single amino acid substitutions. Interface residues are highlighted in red. (H) Heatmaps of inferred ΔΔ*G*_f_ and ΔΔ*G*_b_ (kcal mol^−1^) for KRAS variants binding K13, K19, and RAF1. (I) KRAS sequence and annotation. Binding interfaces: residues within 5 Å of RAF1 or K13; nucleotide-binding pocket: residues within 5 Å of nucleotide or Mg^2+^; core: residues with relative solvent-accessible surface area < 0.25 (AlphaFold-predicted monomer). Secondary structure: P-loop, 10–17; switch-I, 25–40; switch-II, 58–76; α1, 15–24; α2, 67–73; α3, 87–104; α4, 127–136; α5, 148–166; β1, 3–9; β2, 38–44; β3, 51–57; β4, 77–84; β5, 109–115; β6, 139–143.

To quantify the effects of all mutations on binding through both interfaces, we constructed KRAS libraries comprising over 80,000 variants, mostly comprising double aa mutants (Fig. S1A). We generated binding landscapes across the full KRAS sequence for binding partners recognizing the two distinct interfaces (Fig. 1E, Fig. S1B–D; interface 1 binders: RAF1-RBD, K55, and K27; interface 2 binders: K13 and K19). The K13 and K19 binding landscapes were newly generated in this study, and new measurements for RAF1-RBD, K55, and K27 were generated using a synthetic library.

Plotting the binding of each variant against its folded abundance shows that many binding defects are due to reduced abundance of folded KRAS (Fig. 1F), consistent with previous results(*23*). We used MoCHI(*39*) to fit a three-state thermodynamic model to the data to infer the underlying causal changes in the Gibbs free energy of folding (ΔΔ*G*_f_, kcal mol^−1^) and binding (ΔΔ*G*_b_, kcal mol^−1^) for each aa substitution (using combined fitness data from this study and our previous study(*23*); see Methods). The model accounts for the non-linear relationships between the observed molecular phenotypes and the Gibbs free energy of folding (Δ*G*_f_) and binding (Δ*G*_b_) and assumes changes in energy combine additively in double mutants (Fig. 1C). The extent to which the inferred energies predict the observations is highly accurate, with a median Pearson correlation coefficient (R = 0.89) between observed and predicted measurements across all experiments, evaluated by ten-fold cross-validation (Fig. S2A–D). As previously(*23*), the free energy measurements also agree very well with independent *in vitro* measurements (Fig. S2E, R = 0.96). In total, we quantified 26,778 binding free energy changes (ΔΔ*G*_b_) for KRAS variants across eight binders (RAF1-RBD, RALGDS-RBD, PIK3CG-RBD, SOS1, and DARPins K55, K27, K13 and K19; median binding free energy changes per interaction = 3,539 (99.61%)) (Fig. 1H, Fig. S2F, G). Additionally, we quantified 3,551 folding free energy changes (ΔΔ*G*_f_) (Fig. 1H). As an indication of the size of this dataset, the number of ΔΔ*G*_b_ measurements reported is nearly four-fold more than the number reported for all proteins in the largest literature-curated database, SKEMPI v2.0(*40*).

For DARPin K13, 343 of 3,549 quantified substitutions increase the binding free energy (ΔΔ*G*_b_ > 0, lower affinity), whereas 1,756 substitutions decrease ΔΔ*G*_b_ (ΔΔ*G*_b_ < 0, higher affinity). For DARPin K19, 413 of 3,529 substitutions increase ΔΔ*G*_b_, while 681 decrease it. In contrast, for RAF1-RBD, 1,107 of 3,550 substitutions increase ΔΔ*G*_b_, and 605 decrease it (Fig. 1H; false discovery rate (FDR) < 0.05, Benjamini-Hochberg).

### K13 and K19 interfaces

The K13 and K19 binding interfaces are highly similar, with 15 KRAS residues contacting both proteins, three residues only contacting K13 and two only contacting K19 (minimum heavy-atom distance < 5 Å; Fig. 2A). Many mutations in the K13- and K19-binding interfaces disrupt binding (Fig. 1G–I), with median ΔΔ*G*_b_ = 0.37 (n = 130) and 0.17 (n = 166) kcal mol^−1^, respectively, compared to median ΔΔ*G*_b_ = 0.07 (n = 243) and 0.07 (n = 518) kcal mol^−1^ for other surface site mutations (Relative Solvent Accessible Surface Area (RSASA) > 0.25). However, in 3/18 (18.75%) and 3/17 (17.65%) contact sites mutations have much stronger effects, with a median ΔΔ*G*_b_ > 1 kcal mol^−1^, identifying these sites as ‘hotspot’ residues (*41*). The three hotspot residues—E91, H94, and H95—are in the center of both interfaces. H95 forms π-π stacking interactions, whereas E91 contributes a hydrogen bond and salt-bridge interactions, and H94 contributes hydrogen bonds with the interaction partners (Fig. 2A, B).

**Figure 2.**
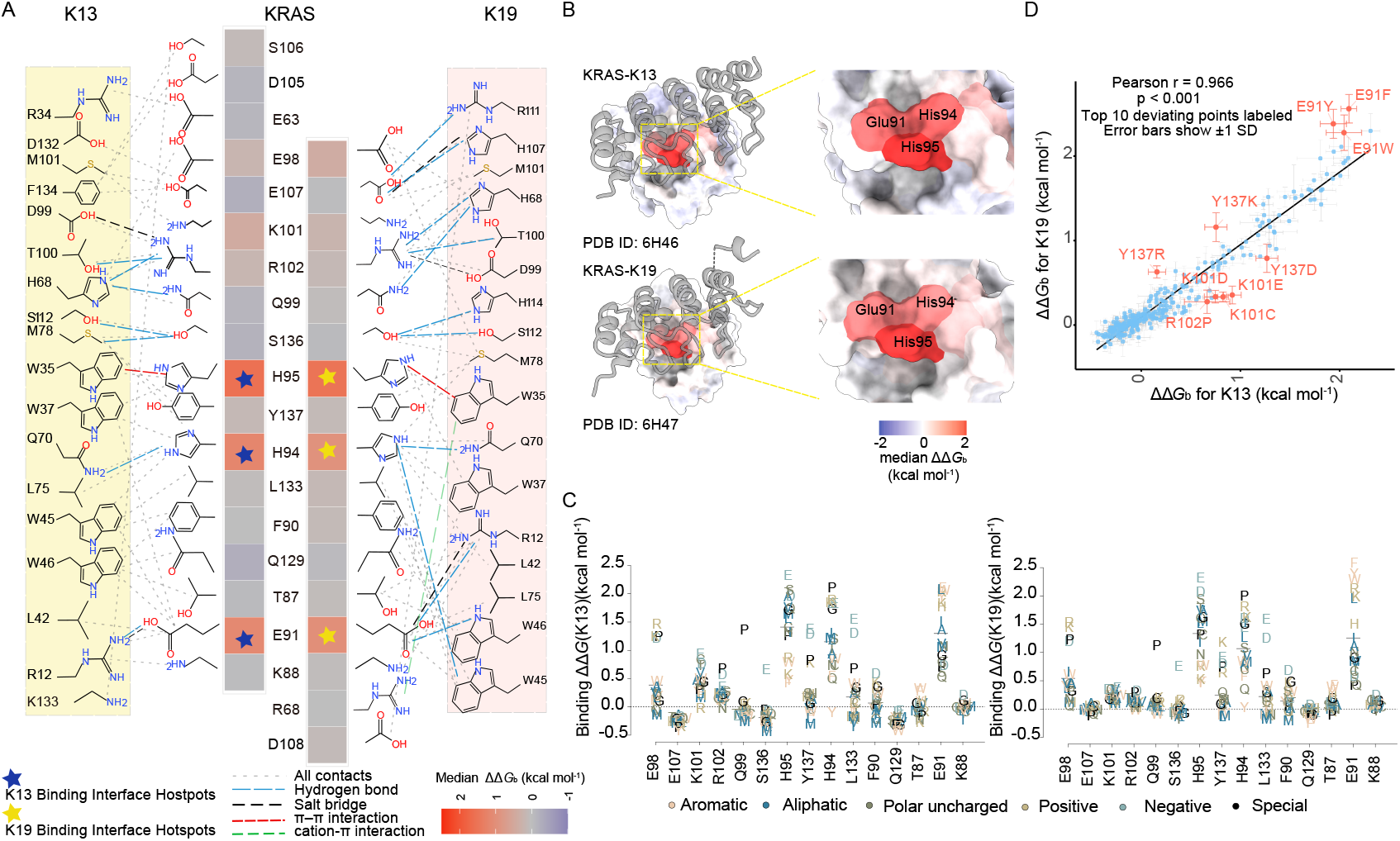
Energy landscapes of the K13 and K19 binding interfaces. (A) Schematic of direct contacts between KRAS and the DARPins, representing the union and unique of interactions with K13 and K19. Interface residues are identified based on contact analysis in UCSF ChimeraX (version 1.12) and structural mapping in Bio3D (RStudio), using a minimum heavy-atom distance cutoff of < 5 Å. Heatmaps show binding free energy changes ΔΔ*G*_b_ (kcal mol^−1^) at interface residues. Blue asterisks denote K13 interface hotspots; yellow asterisks denote K19 interface hotspots. (B) KRAS 3D structures in complex with K13 (top) and K19 (bottom). Residues are colored according to the median ΔΔ*G*_b_ (kcal mol^−1^) of mutations at that position. K13 and K19 are shown in gray ribbons. Magnified views highlight interface hotspots. (C) Violin plots of ΔΔ*G*_b_ (kcal mol^−1^) for mutations at each residue within the union of the K13 and K19 binding interfaces. (D) Correlation of mutational effects on K13 versus K19 binding for residues within the union of the K13 and K19 binding interfaces. Data points corresponding to the ten largest outliers are highlighted in red. Error bars indicate ±1 s.d.; points with large residuals that cannot be explained by measurement error are likely true biologic. Pearson’s R = 0.966, *p* < 10^−16^.

Most mutations in interface 2 have highly correlated ΔΔ*G*_b_ values for binding to K13 and K19 (Pearson’s R = 0.966, *p* < 10^−16^), but some mutations alter the binding specificity, with mutations K101D/E/C, R102P, and Y137D more strongly affecting binding to K13 and mutations Y137R/K and E91Y/F/W more strongly affecting binding to K19 (Fig. 2C, D). Structural analysis suggests the differential effects arise from distinct interactions at the K13 and K19 interfaces (Table S4). For example, the mutation E91F exhibits one of the strongest differential effects and E91 forms a more connected interaction hub with K19 than with K13 (Fig. 2A).

### Comparative allosteric landscapes

Next, we focused on the allosteric regulation of the two structurally-distinct binding interfaces. Mutations outside of a binding interface that alter the binding energy must, by definition, have an indirect—i.e. allosteric—mechanism. The changes in binding energy (ΔΔ*G*_b_, kcal mol^−1^) therefore quantify indirect energetic coupling between each site in a protein and a ligand.

In total, 378 mutations allosterically alter the binding energy for RAF1 with ΔΔ*G*_b_ greater than the mean of the weighted mean absolute binding free energy changes of substitutions at binding interface residues across the eight binders (|ΔΔ*G*_b_| > 0.40 kcal mol^−1^, FDR < 0.05; Fig. 3A). Most of these mutations inhibit binding (n = 357 (94.4%)). Consistent with previous data(*23*), these mutations are strongly enriched in the nucleotide-binding pocket (n = 168, OR = 7.95, *p* < 10^−16^) and also in residues close to the binding interface (127 mutations in the second shell residues relative to the binding interface, OR = 5.42, *p* < 10^−16^; Fig. 3A, Fig. S3A). Allosteric mutations are also enriched in the central β-sheet of KRAS (n = 120, OR = 1.89, *p* = 2.95 × 10^−7^; Fig. 3A, Fig. S3A), with binding free energy changes progressively decreasing in each subsequent strand of the sheet (Fig. S3B).

**Figure 3.**
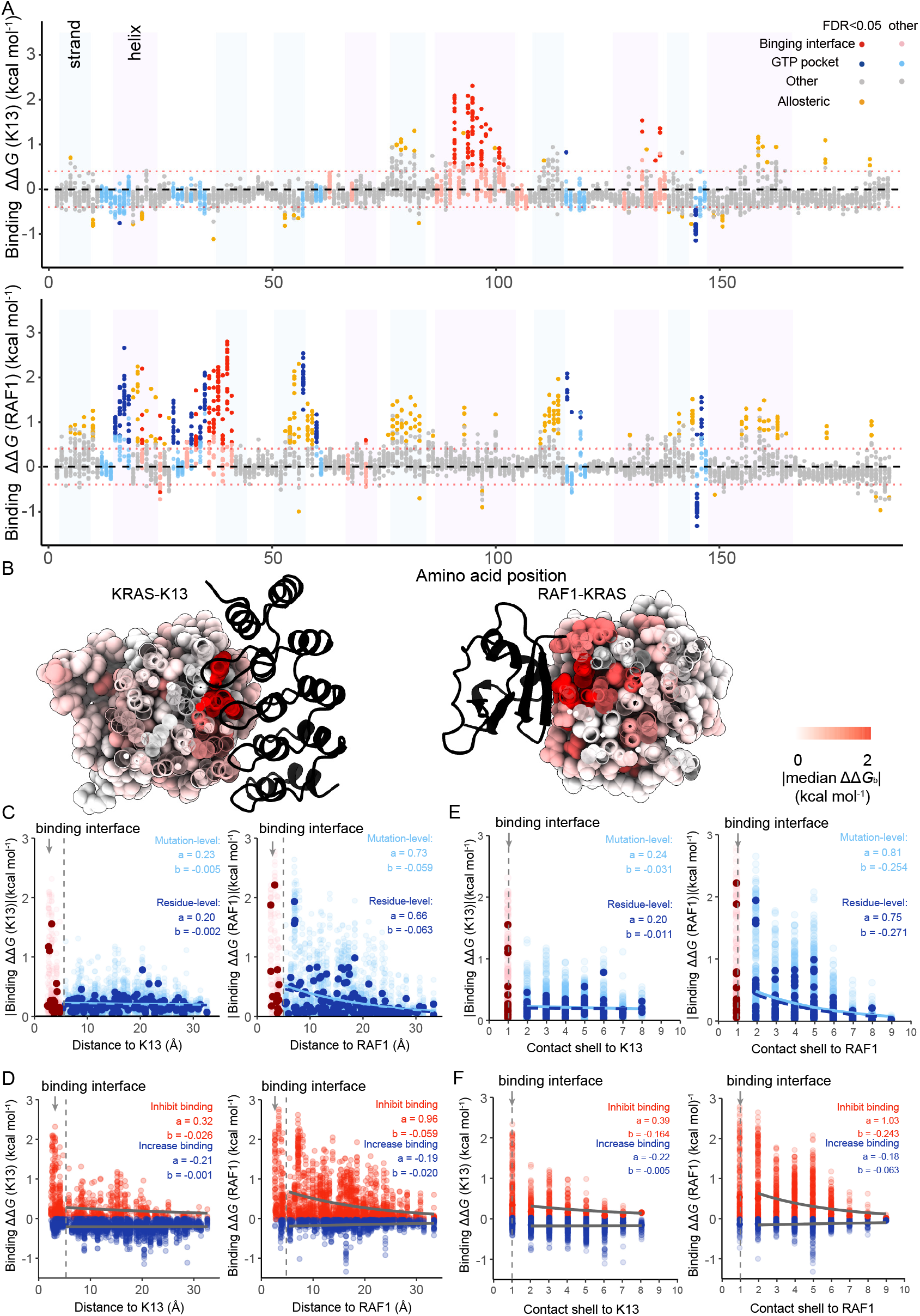
The allosteric landscapes of two KRAS interfaces. (A) Manhattan plot of binding free energy changes ΔΔ*G*_b_ (kcal mol^−1^) for all single amino acid substitutions. Points are colored by residue position; significance is assessed using a two-sided z-test with FDR < 0.05 and |ΔΔ*G*_b_| > 0.40 kcal mol^−1^. (B) Structural visualization of the decay of |ΔΔ*G*_b_| (kcal mol^−1^) with increasing distance from the ligand. KRAS bound to the indicated ligand is shown, colored by median absolute ΔΔ*G*_b_ (kcal mol^−1^) values. (C) Relationship between absolute ΔΔ*G*_b_ (kcal mol^−1^) of K13 and RAF1 and the minimum distance from side-chain heavy atoms to the ligands. Exponential decay fits (*y* = *a* · *e^bx^*) are applied to both mutation-level |ΔΔ*G*_b_| (kcal mol^−1^) values and residue-level median |ΔΔ*G*_b_| (kcal mol^−1^) values. (D) Relationship between ΔΔ*G*_b_ (kcal mol^−1^) of K13 and RAF1 mutations and the minimum distance from side-chain heavy atoms to the ligand. Mutations are stratified into inhibit (ΔΔ*G*_b_ > 0) and increase (ΔΔ*G*_b_ < 0) groups. Exponential decay models (*y* = *a* · *e^bx^*) are independently fitted to mutation-level ΔΔ*G*_b_ values for each group. (E) Relationship between absolute ΔΔ*G*_b_ (kcal mol^−1^) of K13 and RAF1 and the contact shell to the ligands. Exponential decay fits (*y* = *a* · *e^bx^*) are applied to both mutation-level ΔΔ*G*_b_ (kcal mol^−1^) values and residue-level median ΔΔ*G*_b_ (kcal mol^−1^) values. (F) Relationship between ΔΔ*G*_b_ (kcal mol^−1^) of K13 and RAF1 and the contact shell to the ligands. Mutations are stratified into inhibit (ΔΔ*G*_b_ > 0) and increase (ΔΔ*G*_b_ < 0) groups. Exponential decay models (*y* = *a* · *e^bx^*) are independently fitted to mutation-level ΔΔ*G*_b_ values for each group.

For K13 and K19, 77 and 59 allosteric mutations have effects greater than the mean of the weighted mean absolute binding free energy changes of substitutions at binding interface residues across the eight binders (both |ΔΔ*G*_b_| > 0.40 kcal mol^−1^, FDR < 0.05; Fig. 3A, Fig. S3C). 43 mutations meet this criteria for both K13 and K19. In contrast to RAF1, many of the mutations increase binding, corresponding to 48 of 77 (62.3%) for K13. The allosteric mutations for K19 are enriched in the nucleotide-binding pocket (16/59; OR = 2.35, *p* = 6.61 × 10^−3^; Fig. S3A, C) with a weaker enrichment for K13 (17/77, OR = 1.76, *p* = 0.046; Fig. 3A, Fig. S3A) and there is little or no enrichment in second-shell residues relative to the binding interface (all: K13: 13/77; OR = 1.32, *p* = 0.40; K19: 8/59; OR = 0.92, *p* = 1.00; inhibitory: K13 n = 6, OR = 0.92, *p* = 1.00; K19 n = 8, OR = 1.07, *p* = 0.83; activating: K13 n = 7, OR = 1.33, *p* = 0.49; K19 n = 0, OR = 0, *p* = 0.10; Fig. 3A–F, Fig. S3A, C–H).

We define major allosteric sites as residues where the mean absolute change in binding free energy upon mutation is equal or greater than the mean of the weighted mean absolute binding free energy changes binding interface mutations across the eight binders (*22*, *23*) (0.4 kcal mol^−1^). Using this definition identifies twenty-one major allosteric sites for RAF1, three for K13 and three for K19 (Fig. S4). For RAF1, major allosteric sites are enriched in the nucleotide-binding pocket (11/21; OR = 12.67, *p* = 2.73 × 10^−6^) and in second-shell residues relative to the binding interface (9/21; OR = 9.53, *p* = 6.72 × 10^−5^; Fig. S4, Fig. 4A; Movie S4). In contrast, only S145 for K13 and K19 is in the nucleotide-binding pocket and G10 for K13 and K19 is in the binding interface second shell (Fig. S4, Fig. 4A).

**Figure 4.**
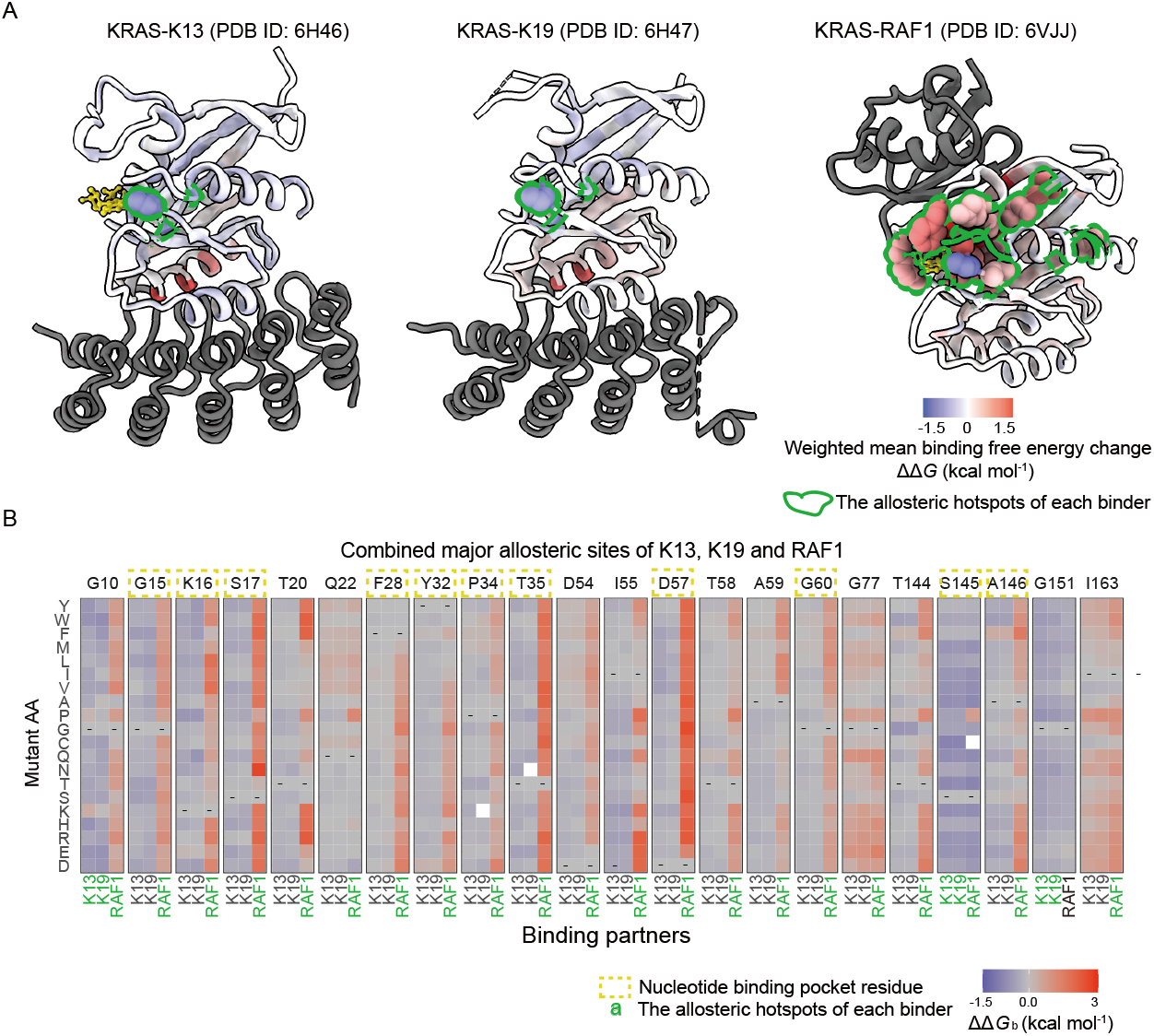
Comparative allosteric regulation of KRAS binding through two interfaces. (A) Three-dimensional structures of KRAS bound to K13, K19, and RAF1 (dark gray) (PDB IDs: 6H46, 6H47, and 6VJJ). Residues are colored using weighted mean free energy values calculated independently for each complex. Green outlines indicate residues identified as allosteric hotspots in each binder. (B) Heatmaps of binding free energy changes in allosteric residues combined by three binding partners (K13, K19, and RAF1). The residues of the nucleotide-binding pocket site are enclosed within the yellow dashed-line border. Green front indicate residues identified as allosteric hotspots in each binder.

We identified 22 residues that were classified as major allosteric sites for at least one of the three interaction partners (Fig. 4B). Among these, S145, located in the loop connecting β-strand 6 and α-helix 5, is a major allosteric site shared by all three binders (RAF1, K13, and K19). S145 contacts the nucleotide through the O6 carbonyl oxygen of the guanine base of GDP or GNP (PDB IDs: 6H46 and 6VJJ) but does not interact with the phosphate groups. Except for S145P, substitutions at this position enhance binding to all three interaction partners (Fig. 4A, B).

### Three modes of allosteric regulation

To further compare allosteric regulation of the two interfaces, we compared the effects of all mutations outside of both interfaces on binding. Plotting the binding free energy changes for the two interface 2 binders, K13 and K19, reveals that they are highly correlated (Fig. 5A; Pearson’s R = 0.886, *p* < 10^−16^). Similarly, comparing the binding free energy changes for RAF1 to those for five additional interface 1 binders(*23*) shows that they are also very well correlated (Fig. 5A, Fig. S5A; median Pearson’s R = 0.725, *p* < 0.001).

**Figure 5.**
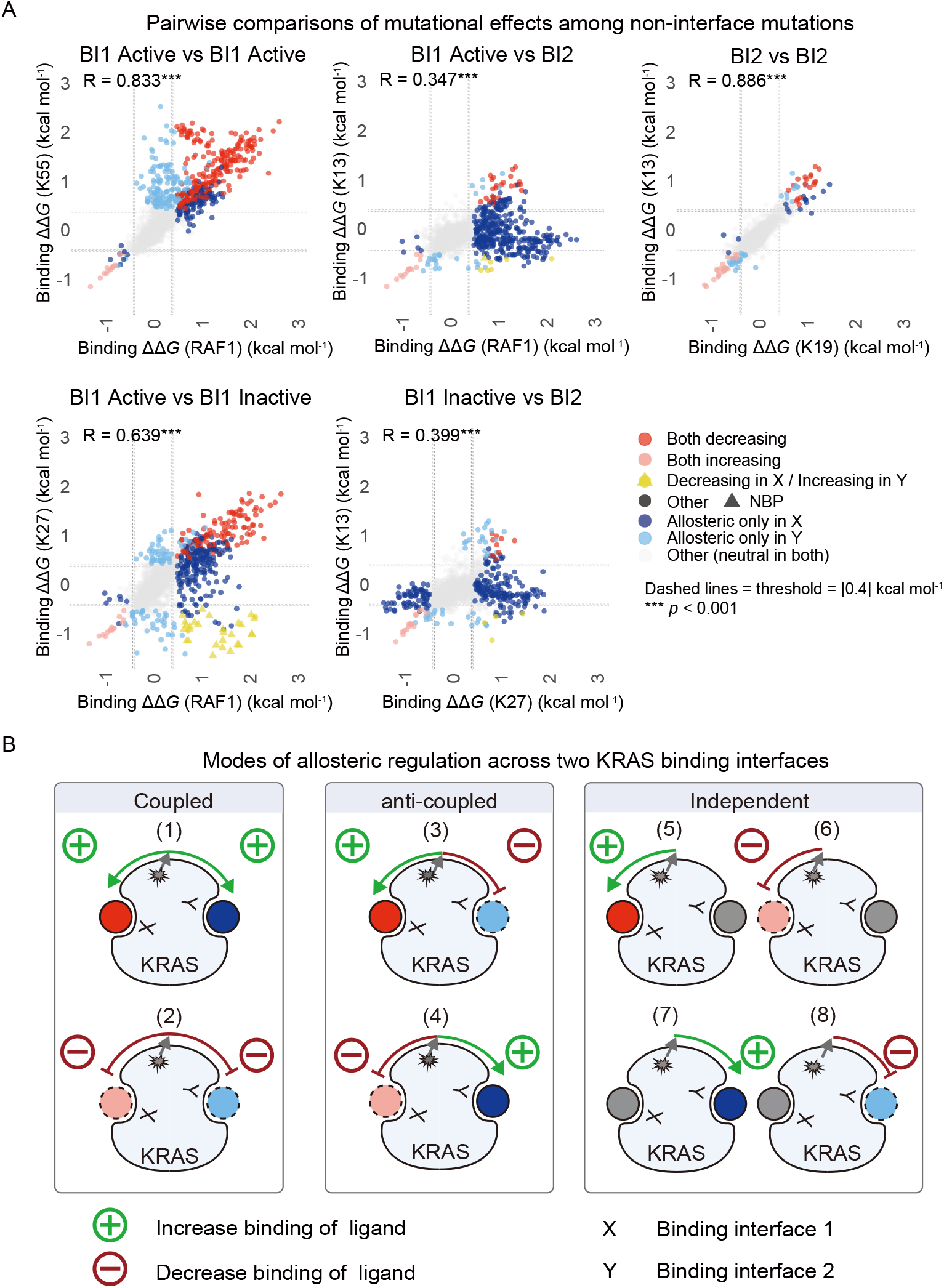
Three forms of allosteric regulation for two KRAS binding interfaces. (A) Pairwise scatter plots of mutational effects for mutations outside the two compared binding interfaces, across BI1-active versus BI1-active, BI1-active versus BI1-inactive, BI1-active versus BI2, BI1-inactive versus BI2, and BI2 versus BI2 conditions. BI1 denotes interface 1 binding, with “active” and “inactive” referring to the corresponding KRAS binding states; BI2 denotes interface 2 binding. Each point represents a mutation. Colors indicate effects exceeding the threshold in both conditions (red, decreasing; pink, increasing), opposite effects exceeding the threshold (yellow), or a threshold-exceeding effect specific to the X-axis (blue) or Y-axis (light blue); grey indicates neutral effects. Effects are considered significant at FDR < 0.05 and a threshold of 0.40 kcal mol^−1^, defined as the mean of the weighted mean |ΔΔ*G*_b_| of substitutions at binding interface residues across all eight binders. (B) Cartoon illustrating three forms of allosteric regulation across two KRAS interfaces. Coupled mutations affect both interfaces in the same direction (either increasing or decreasing binding), whereas anti-coupled mutations have opposite effects on the two interfaces. Independent mutations selectively affect binding at one interface with minimal or no effect on the other.

In contrast, comparing the changes in binding energy for six different interface 1 binders (RAF1-RBD, RALGDS-RBD, PIK3CG-RBD, SOS1, and DARPins K55, K27) to those for the two interface 2 binders (DARPins K13 and K19) reveals an unexpected and more complicated pattern, with four different behaviors (Fig. 5A, B, Fig. S5A). First, a set of mutations has correlated effects on interface 1 and interface 2, either inhibiting or activating both. For RAF1 vs. K13 (Fig. 5A, B), this set is twenty-three mutations in fifteen residues that inhibit binding and fourteen mutations in three residues that enhance binding to both proteins (inhibit binding: ΔΔ*G*_b_ (RAF1) > 0.40 kcal mol^−1^, ΔΔ*G*_b_ (K13) > 0.40 kcal mol^−1^; enhance binding: ΔΔ*G*_b_ (RAF1) < −0.40 kcal mol^−1^, ΔΔ*G*_b_ (K13) < −0.40 kcal mol^−1^, FDR < 0.05; ΔΔ*G*_b_ greater than the mean of the weighted mean absolute binding free energy changes of substitutions in binding interface residues across the eight binders). Moreover, the effects of these thirty-seven mutations are well correlated for the two proteins (Pearson’s R = 0.98, *p* < 10^−16^). We refer to these as ‘coupled’ allosteric mutations as their effects on binding are similar for the two interfaces.

Second, a group of seven mutations in five residues inhibits binding to RAF1 but increases binding to K13 (ΔΔ*G*_b_ (RAF1) > 0.40 kcal mol^−1^, ΔΔ*G*_b_ (K13) < −0.40 kcal mol^−1^, FDR < 0.05; ΔΔ*G*_b_ greater than the mean of the weighted mean absolute binding free energy changes of substitutions in binding interface residues across the eight binders). We refer to these mutations as inversely-coupled or anti-coupled allostery, as the effects are opposite on the two interfaces. Interestingly, the mutations with anti-correlated effects on interface 1 and interface 2 binding are not the same as those with anti-correlated effects on active state (RAF1) vs. inactive state (K27) interface 1 binders (Fig. S5C–E), suggesting that their mechanism of action is not (only) a shift in the equilibrium between the KRAS active (GTP-bound) and inactive (GDP-bound) states.

Third, 325 mutations inhibit and 5 mutations promote binding to RAF1 but have little effect on binding to K13 (|ΔΔ*G*_b_ (RAF1)| > 0.40 kcal mol^−1^, |ΔΔ*G*_b_ (K13)| < 0.40 kcal mol^−1^, FDR < 0.05). Fourth, six mutations inhibit and nineteen promote binding to K13 but have little effect on RAF1 binding (|ΔΔ*G*_b_ (RAF1)| < 0.40 kcal mol^−1^, |ΔΔ*G*_b_ (K13)| > 0.40 kcal mol^−1^, FDR < 0.05). We refer to these last two sets of mutations as independent allostery, as the mutations only strongly affect binding to one of the two interfaces. Across all interface 1 vs. interface 2 binder comparisons, there are a larger number of independent allosteric mutations for interface 1 than for interface 2 (Fig. S5B).

### Coupled, inversely-coupled and independent allostery across the KRAS structure

To investigate the spatial organization of the different classes of allosteric mutations, we mapped mutations exhibiting coupled, inversely-coupled, and independent allostery onto the 3D structures of KRAS in complex with RAF1 and K13 (Fig. 6A, PDB IDs: 6VJJ and 6H46; Movie S5). Both coupled and inversely-coupled mutations are enriched in the protein core (Fig. 6A, B; Fig. S6A; coupled OR = 4.16, *p* = 1.69 × 10^−4^; inversely-coupled OR = ∞, *p* = 5.01 × 10^−3^). Within the protein core, coupled mutations are enriched in β-strand 4 and its surrounding region (OR = 7.05, *p* = 1.30 × 10^−5^), whereas inversely-coupled mutations are enriched in β-strand 3 (OR = 28.32, *p* = 1.35 × 10^−4^). β-strand 3 is positioned closer to the effector-binding interface, whereas β-strand 4 is located deeper within the central β-sheet in the core (Fig. 6A). Thus, although both mutation classes cluster within the central β-sheet region, they are enriched in distinct regions within it. However, this spatial segregation is not absolute, with 17 of 18 residues (94.4%) harboring coupled mutations and all 5 residues (100%) harboring inversely-coupled mutations also contain mutations from other allosteric classes, indicating that individual residues can encode multiple modes of allosteric regulation depending on the specific amino acid substitution (Fig. 6A). For example, substitutions at S145 produce coupled, inversely-coupled and K13-specific allosteric effects.

**Figure 6.**
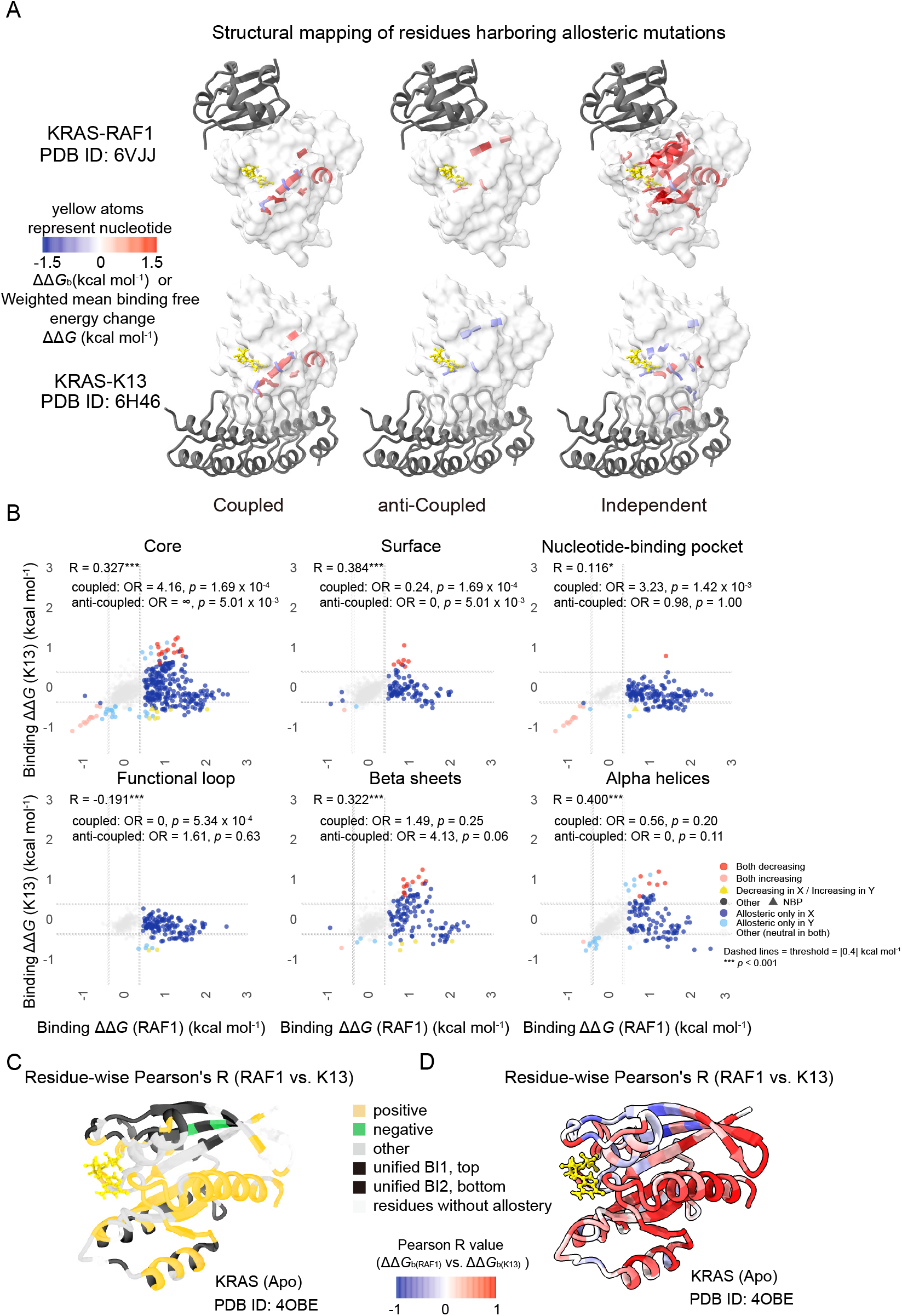
Coupled, anti-coupled and independent allostery across the KRAS structure. (A) Residues harboring coupled, anti-coupled, or independent allosteric mutations from the RAF1 vs. K13 comparison are mapped onto KRAS structures in complex with RAF1 and K13 (PDB IDs: 6VJJ and 6H46). Only residues containing at least one mutation classified into the corresponding allosteric category are colored. When multiple mutations are present at the same site, weighted mean ΔΔ*G*_b_ values are used. (B) Pairwise scatter plot of mutational effects for mutations outside the RAF1 and K13 binding interfaces, comparing RAF1 versus K13 binding. Mutations are stratified by structural regions of KRAS, including the protein core (RSASA < 0.25), solvent-exposed surface (RSASA > 0.25), nucleotide-binding pocket (NBP), functional loops (switch I, switch II, and P-loop), β-strands 1–6, and α-helices 1–5. Each point represents a mutation. Colors are as defined in Fig. 5A. (C) Three-dimensional structure of KRAS showing the four groups of residues defined in Fig. S6C. Orange, positive correlation residues supporting allostery (*p* < 0.05); green, negative correlation residues supporting allostery (*p* < 0.05); light grey, other correlation residues supporting allostery (*p* > 0.05); black, residues in the union of BI1 (RAF1/K55/K27; top) or BI2 (K13/K19; bottom); white, residues without allostery (residues supporting allostery indicate that the residue has a significant allosteric mutation (defined in Fig. 5A). (D) Three-dimensional structure of KRAS (PDB ID: 4OBE). Each residue is colored according to its correlation coefficient (Pearson’s R value) in Fig. S6C. The color scale indicates the range of R values.

Mutations that only strongly allosterically affect RAF1 binding are enriched close to the RAF1 interface (OR = 6.12, *p* < 10^−16^ in the second shell relative to binding interface) and are also enriched in the nucleotide-binding pocket (OR = 7.28, *p* < 10^−16^) (Fig. 6A, Fig. S6B). Similarly, mutations that only strongly allosterically affect K13 binding are enriched near the K13-binding interface (OR = 2.43, *p* = 0.07 in the second shell relative to binding interface) but they are not enriched within the nucleotide-binding pocket (OR = 0.51; *p* = 0.57) (Fig. 6A, Fig. S6B). Instead, K13-specific mutations are enriched in α-helical regions (OR = 3.69, *p* = 1.40 × 10^−3^), although they are broadly distributed across multiple α-helices (Fig. 6A, Fig. S6B).

The above analyses classify individual mutations according to their effects on the two binding interfaces binders (RAF1 vs. K13). We next quantified the correlation between ΔΔ*G*_b_ values for RAF1 and K13 across the 19 mutations at each residue (Fig. S6C, D). Visualizing these correlation coefficients on the 3D structure of KRAS (Fig. 6C, D; Movie S6, 7) reveals that residues where mutations have positively correlated effects on RAF1 and K13 binding are predominantly located in β-strand 4 (7/8 residues, OR = 5.05, *p* = 0.145), consistent with the enrichment of coupled allosteric mutations in this strand (Fig. 6, Fig. S6A). Only two residues exhibit significant negative correlation (L53 and I55, *p* < 0.05) but residues with indicative negative correlation coefficients form a structural cluster centered on the nucleotide-binding pocket (Fig. 6C, D, Fig. S6C). These residues are enriched in the nucleotide-binding pocket (OR = 16.93, *p* = 4.13 × 10^−6^), with additional enrichment in the γ-phosphate-contacting residues (OR = 14.08, *p* = 7.30 × 10^−3^) and β-strand 3 (OR = 10.50, *p* = 0.012). The enrichment in β-strand 3 is consistent with the enrichment of inversely-coupled mutations in this region at the mutation level (Fig. 6A, B, Fig. S6A).

### Multimodal allosteric regulation from the KRAS surface and pockets

For protein regulation and therapeutic targeting, solvent-exposed surface sites are of particular interest(*5*). We therefore examined the effects of surface mutations (RSASA > 0.25) on the binding to RAF1 and K13 (Fig. 7). In total, 103 surface-accessible mutations exhibit allosteric effects (|ΔΔ*G*_b_ (RAF1)| > 0.40 kcal mol^−1^, |ΔΔ*G*_b_ (K13)| > 0.40 kcal mol^−1^, FDR < 0.05), including eight coupled mutations, one K13-specific mutation, and ninety-four RAF1-specific mutations. Coupled mutations are distributed across multiple surface regions of KRAS (Table S5) whereas RAF1-specific mutations are enriched in the effector lobe (eighty-four mutations in eleven residues, OR = 4.62, *p* = 3.0 × 10^−6^) and nucleotide-binding pocket region of KRAS (seventy-two mutations in seven residues, OR = 6.34, *p* < 10^−16^; Table S5).

**Figure 7.**
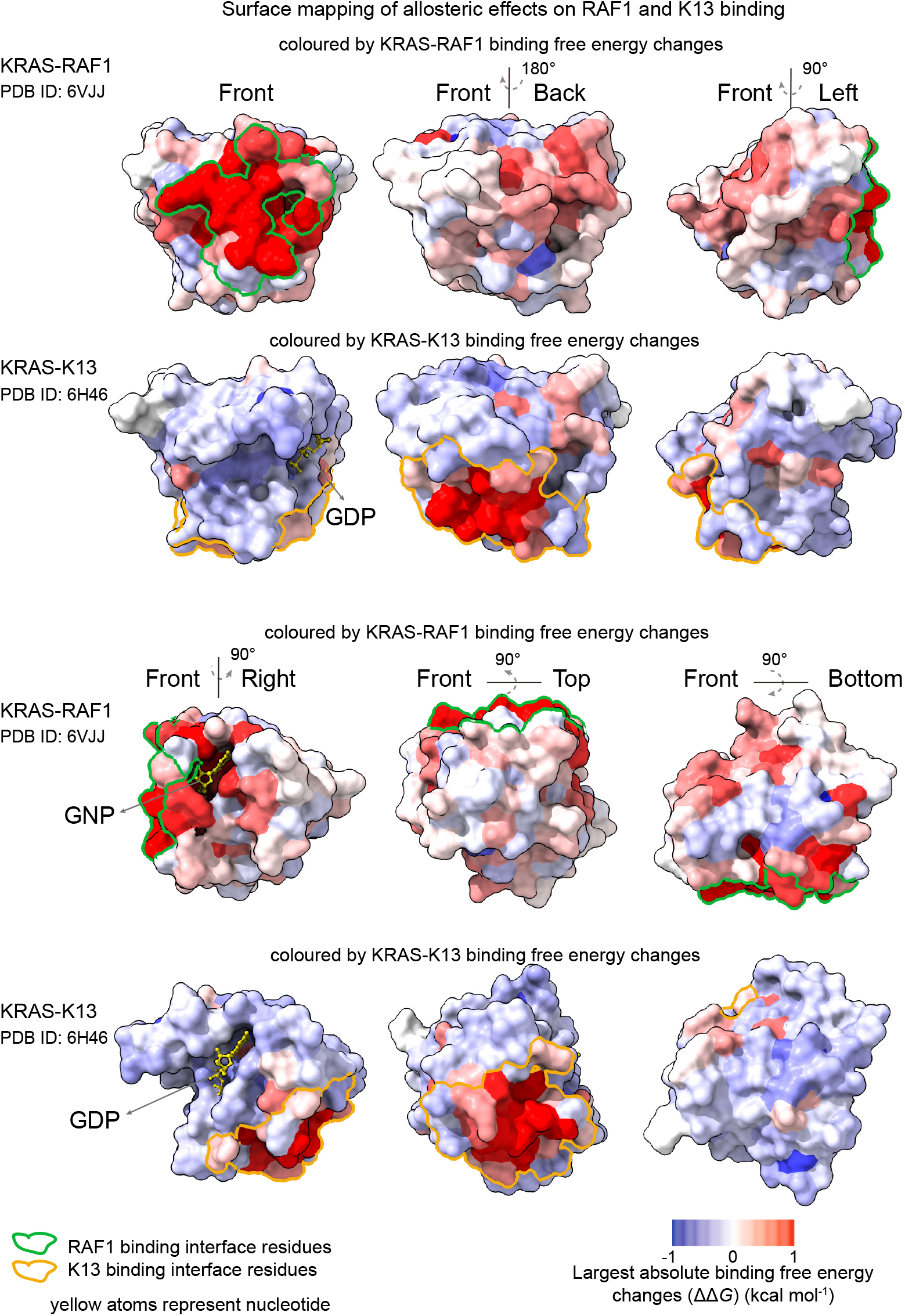
Multimodal allosteric regulation from the KRAS surface and pockets. Structural alignment of KRAS bound to RAF1 and K13 (PDB IDs: 6VJJ and 6H46) with KRAS surface in “front” (relative to interface 1), “back”, “left”, “right”, “top”, and “bottom” views, coloured according to the signed ΔΔ*G* values of mutations with the largest absolute binding free energy changes for RAF1 and K13 at each residue, the colour scale is capped at −1 and +1 kcal mol^−1^; values beyond this range are shown at the corresponding colour-scale limits. The “back”, “left”, “right”, “top”, and “bottom” views are generated by rotation relative to the “front” view, arrows indicate the direction of rotation. Green outlined surface represents interface 1; Orange outlined surface represents interface 2. Surface accessibility is determined using the full-length AlphaFold model of KRAS (residues 1–188; RSASA > 0.25). The mutations are located in the HVR, which is absent from the crystal structure used for visualization and therefore are not displayed.

We next considered four well-characterized non-nucleotide-binding pockets of KRAS(*42*). All four pockets harbor allosteric mutations affecting RAF1 and/or K13 binding (FDR < 0.05; Fig. S7; Table S5). Pocket 1 (the switch-I/II pocket) is located between the central β-sheet and α2 helix behind switch II and is the binding site for multiple pre-clinical small-molecule inhibitors(*43, 44*). This pocket contains thirty-three allosteric mutations, including mutations with coupled, inversely-coupled and RAF1-specific effects. Pocket 2 (the switch-II pocket) is located between switch II and the α3 helix and is bound by clinically approved KRAS inhibitors(*15*). Several residues in this pocket overlap the K13 binding interface (E63, H95, and Q99) and fifty-four mutations in this pocket exhibit inversely-coupled, RAF1-specific, and K13-specific effects. Pocket 3 is located in the C-terminal lobe of KRAS, furthest from the RAF1 binding interface and partially overlaps the K13 binding interface(*42, 44*). This pocket contains twenty-four allosteric mutations, with coupled, RAF1-specific, and K13-specific effects. Pocket 4 is positioned behind the effector-binding loop(*42*) and contains ninety-four RAF1-specific allosteric mutations.

Thus, perturbations originating from the protein surface and pockets of KRAS can exert correlated, anticorrelated, and independent effects on binding to the two interfaces, suggesting the potential for correlated, anticorrelated and independent physiological or therapeutic control of two interfaces in a single protein.

## Discussion

Allostery allows the activities of proteins to be controlled by events outside of their active sites. Using mutations as perturbations it is possible to quantify allosteric communication between all sites in a protein and an active site. To data, however, complete maps of allosteric mutations have only been constructed for a single target site in each protein(*11*, *22–25*). Many proteins, however, have multiple functional sites. How the allosteric regulation of two distinct sites maps throughout a protein structure is unknown.

Here we have directly addressed this question by constructing complete allosteric maps for binding to two spatially distant sites in the same protein, the oncoprotein KRAS. Quantifying thousands of binding energy changes allowed us to build comprehensive allosteric maps for both binding interfaces, quantifying the effects of 19 different perturbations at every site in the protein. Very strikingly, these maps reveal that three distinct forms of allosteric regulation co-exist within the same protein, with different sets of mutations allosterically regulating both interfaces in a correlated manner, having anticorrelated effects on the two interfaces, or only affecting one of the two interfaces. We refer to these three allosteric modes as coupled, anti-coupled and independent allostery.

The co-existence of coupled, anti-coupled and independent allostery has important implications for our understanding of protein energy landscapes. Allostery is often conceptualized as a shift between two alternative conformational or energetic states(*45*, *46*). However, such a model predicts mutations should only have positively or negatively correlated effects on binding, depending upon whether the interaction partners bind the same or alternative states. The coexistence of coupled, anti-coupled and independent allostery within KRAS, a small single-domain protein, is therefore unexpected. Such multimodal regulation is difficult to reconcile with classical models of allostery and instead suggests that multiple overlapping allosteric networks coexist within the same protein. In KRAS there must be at least four distinct allosteric networks (or mechanisms) in the protein: one that co-regulates both interfaces, one that has opposite effects on them, one that only affects interface 1, and one that only affects interface 2. The distinct but overlapping spatial organization of allosteric mutations in each class further supports this. Mechanistically, changes in either conformation or dynamics could underlie each allosteric network, with this potentially differing both across proteins and across networks(*5*, *47*, *48*). Alternatively, a complex ensemble of energy states might exist, with each state favoring binding to interface 1, interface 2, both, or neither and mutations shifting the distribution between these many states. We note that multimodal allosteric regulation is also observed for overlapping interfaces, with the existence of both coupled and anti-coupled allosteric mutations for the interface 1 binders RAF1 and K27(*23*) (Fig. 6b in Weng *et al*).

Our results also have implications for cellular regulation and evolution. Coupled, anti-coupled, and independent allosteric control of two protein functions would provide flexibility to cellular regulation, allowing, for example, different post-translational modifications to inhibit two functions, inhibit only one function, or inhibit one but activate another. During evolution, a new binding partner for a protein might be positively or negatively coupled to existing allosteric regulation or bind independent of existing regulation.

Allosteric drugs often have multiple desirable characteristics, including increased selectivity, reduced toxicity, different sets of resistance mutations to orthosteric drugs, and the ability to modulate ‘undruggable’ proteins that lack an active site that can be targeted with small molecule(*5*, *17*, *49*). The co-existence of multiple modes of allosteric co-regulation in a protein suggests the potential for new classes of allosteric therapeutics that not only modulate a target but also tune its functional outputs. Such allosteric functional modulators would be conceptually similar to biased receptor agonists(*50*). We speculate that targeting different molecules to the same allosteric site or targeting different allosteric sites may allow the development of novel therapeutics that specifically modulate only one protein function, multiple functions, or inhibit one but activate another.

## Data availability

All sequencing data have been deposited in the European Nucleotide Archive (ENA) at EMBL-EBI under accession number PRJEB123262.

## Code availability

Source code for fitting thermodynamic models (MoCHI) is available at https://github.com/lehner-lab/MoCHI. Source code for fitness analyse is available at https://github.com/lehner-lab/DiMSum. Source code for all downstream analyses and to reproduce all figures described here is available at https://github.com/weng-lab-ustc/Multimodal-allostery.

## Acknowledgments

This work was supported by grants from the National Natural Science Foundation of China (32571676 to C.W.), Wellcome (220540/Z/20/A), the European Research Council (ERC, Advanced Grant 883742), the Spanish Ministry of Science and Innovation (PID2023-146685NB-I00, EMBL Partnership, Severo Ochoa Centre of Excellence), AGAUR (2021 SGR 01226), and the CERCA Programme/Generalitat de Catalunya. We thank all members of the Lehner and Weng labs, and Professor Long Dong for helpful discussions and suggestions.

## Author contributions

B.L. and C.W. conceived the project. Y.M., B.L. and C.W. designed the experiments and analyses. Y.M., Q.J., and C.W. performed the experiments. Y.M. and C.W. performed the data analysis. Y.M., Q.J., B.L., and C.W. wrote the manuscript.

## Supplementary Information

### Materials and Methods

#### Media and Buffers

The following media and buffers were used in this study and prepared as described below:

Luria-Bertani (LB) medium was formulated as follows: per liter, 10 g of Bacto-tryptone, 5 g of yeast extract, and 10 g of NaCl; autoclaved at 121 °C for 20 minutes.

Yeast Peptone Dextrose Adenine (YPDA) medium contained per liter 20 g of glucose, 20 g of peptone, 10 g of yeast extract, and 40 mg of adenine hemisulfate; autoclaved at 115 °C for 30 minutes.

Sorbitol medium (SORB) was composed of 1 M sorbitol, 100 mM lithium acetate, 10 mM Tris-HCl (pH 8.0), and 1 mM EDTA; sterilized by filtration through a 0.2 μm nylon membrane (Merck).

Plate mixture was formulated with 40% PEG3350, 100 mM lithium acetate, 10 mM Tris-HCl (pH 8.0), and 1 mM EDTA (pH 8.0); sterilized by membrane filtration.

Recovery medium utilized yeast glucose medium (containing 20 g L^−1^ glucose, 20 g L^−1^ peptone, and 10 g L^−1^ yeast extract) supplemented with 0.5 M sorbitol; sterilized by filtration.

Synthetic Complete medium (without uracil, SC-URA) was formulated per liter with 6.7 g of yeast nitrogen base without amino acids, 20 g of glucose, and 0.77 g of supplement mixture without uracil; sterilized by membrane filtration.

SC-URA/MET/ADE medium was formulated without uracil, methionine, and adenine, with the following composition per liter: 6.7 g of yeast nitrogen base, 20 g of glucose, and 0.74 g of supplement mixture lacking uracil, adenine, and methionine; similarly sterilized by membrane filtration.

The medium used for competition assays was based on SC-URA/MET/ADE, supplemented with 200 μg ml^-1^ methotrexate (BioShop Canada) and 2% DMSO. DNA extraction buffer was composed of 2% Triton X-100, 1% SDS, 100 mM NaCl, 10 mM Tris-HCl (pH 8), and 1 mM EDTA (pH 8).

#### Plasmid Construction

Two universal plasmids for detecting protein-protein interactions or protein abundance via BindingPCA or AbundancePCA, respectively, were constructed: the BindingPCA plasmid (pGJJ161) and the AbundancePCA plasmid (pGJJ162). These plasmids were derived from previously developed backbones—pGJJ001 (BindingPCA) and pGJJ045 (AbundancePCA). A key modification involved relocating the C-terminal (GGGGS)₄ linker to the N-terminus of the DHFR3 fragment, enabling fusion of target proteins to the N-terminus of DHFR3 for use in either assay. This study utilized one KRAS AbundancePCA plasmid, two KRAS BindingPCA plasmids, and one KRAS mutant library plasmid.

To construct the KRAS AbundancePCA plasmid (pGJJ271), the full-length KRAS sequence (188 amino acids) was amplified from a plasmid provided by the L. Serrano laboratory using primers oGJJ231 and oGJJ232, which introduced HindIII and NheI restriction sites. The PCR product was digested with HindIII and NheI and ligated into the similarly digested pGJJ162 vector using T4 DNA ligase (NEB).

For the KRAS BindingPCA plasmids, a universal acceptor plasmid (pGJJ317) was first generated by cloning the HindIII/NheI-digested full-length KRAS insert into the BindingPCA backbone. Two specific BindingPCA plasmids were then constructed by cloning PCR-amplified binding protein sequences into the BamHI/SpeI sites of pGJJ317. The DARPin K13 BindingPCA plasmid (pGJJ683) was generated by amplifying the DARPin K13 (residues 1–156) sequence from plasmid pCASP-SptP120-K13-HilA (Addgene) using primers oWCC324 and oWCC325. Similarly, the DARPin K19 BindingPCA plasmid (pGJJ684) was constructed by amplifying the DARPin K19 (residues 1–157) sequence from plasmid pCASP-SptP120-K19-HilA using primers oWCC326 and oWCC327.

The KRAS mutant library plasmid (pGJJ380) was constructed in several steps. First, an intermediate plasmid, pGJJ191, containing a streptomycin resistance cassette, was assembled via Gibson assembly from two PCR fragments: one containing the origin of replication with introduced AvrII and HindIII sites (amplified with primers oGJJ308/oGJJ309) and the other containing the streptomycin resistance gene (amplified with primers oGJJ310/oGJJ311). The KRAS sequence was then excised from the AbundancePCA plasmid using AvrII and HindIII and cloned into the corresponding sites of pGJJ191. Finally, a BbvCI restriction site was introduced using primers oWCC51 and oWCC52 to facilitate subsequent library construction.

#### Mutant Library Construction

To enable full-length sequencing on the Illumina PE150 NextSeq platform, the KRAS gene was divided into three blocks. In this study, block 1 was mutagenized using a plasmid-based one-pot saturation nicking mutagenesis method(*1*), while blocks 2 and 3 were constructed using two distinct approaches: the one-pot saturation nicking mutagenesis method and synthetic mutagenesis.

For block 1 mutagenesis, an initial round of nicking mutagenesis was performed using an equimolar mixture of degenerate KRAS primers (see Table S1). This step served two purposes: (1) to generate a pool of random single mutants, which were used as templates for subsequent rounds of mutagenesis (individual clones were randomly selected and validated by Sanger sequencing); (2) to assess the amplification bias of the degenerate primers at each position, allowing for subsequent bias compensation in the constructed shallow double-mutant libraries.

To construct the three final KRAS libraries, a single round of nicking mutagenesis was performed using an equimolar pool of single mutants for each block and the wild-type sequence as plasmid templates. Mutants were selected based on their varying binding affinities for RAF1(*2*, *3*) to ensure the libraries encompassed a range of affinities. The selected mutants were: block 1: T2K, V14S, L6H, E37G, Y40A, D38C, L19P, Q61L, E63V; block 2: I84L, F82S, L113F, Y71F, K101R, A66P, M72G, F78W, E63V, V112N; block 3: K176C, R149V, L133A, Y137K, L159A, A146F. Additionally, mutants of interest (G12C, G12D, G12V, S17N, and T35S) were included in block 1. To compensate for extreme positional bias, the mixing ratio of each mutagenic primer in the pool was inversely proportional to the average read count per position from the first-round nicking libraries.

For the synthetic mutagenesis of blocks 2 and 3, NNK saturation mutagenesis was performed at every amino acid position within each block, using background mutations N116 (block 2) and S145 (block 3) as scaffolds. Each NNK (N = A/T/C/G; K = T/G) degenerate codon encodes 32 possible codons, yielding a final library of 64,512 unique nucleotide sequences.

For each synthesized pool, primers were designed to amplify the entire pool, and mutant oligonucleotides were introduced via Gibson assembly. Library midi-preparations were digested with HindIII and NheI restriction enzymes, and inserts containing the mutant proteins were gel-purified using the MinElute Gel Extraction Kit (QIAGEN). These were then cloned into the AbundancePCA and BindingPCA plasmid backbones via temperature-cycled ligation. Both AbundancePCA and BindingPCA plasmid backbones were digested with HindIII and NheI and purified using the QIAquick Gel Extraction Kit (QIAGEN). Assembly of the AbundancePCA and BindingPCA libraries was completed by overnight temperature-cycled ligation with T4 DNA ligase (New England Biolabs) according to the manufacturer’s protocol, using 67 fmol of backbone and 200 fmol of insert in a 33.3 µL reaction. Ligations were desalted by dialysis using membrane filters for 1 hour and subsequently concentrated 3.3-fold using a SpeedVac concentrator (HENGNOU).

According to the manufacturer’s protocol, all concentrated assembled libraries were transformed into E. coli DH5α or DH10β (WEIDI) high-efficiency electrocompetent cells. Cells were recovered in SOC medium for 60 minutes and then transferred into 20 mL of LB medium containing ampicillin (final concentration 25 µg mL^−1^). The estimated total number of transformants for each library is provided in Table S1. Following overnight incubation, saturated E. coli cultures for each library were harvested, and the plasmid libraries were extracted using the QIAfilter Plasmid Midi Kit (QIAGEN).

#### Large-Scale Transformation and Competition Assay

Variant libraries were transformed in triplicate with a minimum coverage of 30-fold (average library coverage ≥ 100×). For each selection assay (comprising 3 blocks × 5 BindingPCA assays + 3 blocks × 1 AbundancePCA assay), three independent pre-cultures of BY4742 were grown overnight at 30 °C in 20 mL of standard YPDA medium. The following morning, cultures were diluted to an OD600 of 0.3 in 175 mL of pre-warmed YPDA and incubated at 30 °C for 4 hours. Cells were then harvested by centrifugation at 3,000 g for 5 minutes, washed with sterile water, and subsequently centrifuged in SORB medium (100 mM lithium acetate, 10 mM Tris-HCl pH 8.0, 1 mM EDTA, 1 M sorbitol). The cell pellet was resuspended in 8.6 mL of SORB and incubated at room temperature for 30 minutes.

Following incubation, 175 μL of 10 mg mL^−1^ denatured salmon sperm DNA (Solarbio) and 3.5 μg of plasmid library were added to each tube of cells. After gentle mixing, 35 mL of plate mixture was added to each tube, followed by an additional 30-minute incubation at room temperature. Then, 3.5 mL of DMSO was added to each tube, and the cells were subjected to heat shock at 42 °C for 20 minutes, with periodic inversion to ensure uniform heat transfer. After heat shock, cells were centrifuged and resuspended in approximately 50 mL of recovery medium, followed by a 1-hour recovery period at 30 °C (tubes were inverted every 15 minutes). Cells were centrifuged again, washed with SC-URA medium, and resuspended in SC-URA (volumes used per library are listed in Table S1). After homogenization by stirring, 10 μL of the suspension was plated on SC-URA plates and incubated at 30 °C for approximately 48 hours to determine transformation efficiency. Independent liquid cultures were grown at 30 °C for about 48 hours until saturation. The number of yeast transformants obtained for each library assay is provided in Table S1.

For each BindingPCA or AbundancePCA assay, growth competition was initiated immediately after yeast transformation. Following the first cycle of plasmid selection post-transformation, a second selection cycle was performed by inoculating SC-URA/MET/ADE medium at a starting OD600 of 0.1 using saturated cultures (volumes per experiment specified in Table S1). Cells were grown at 30 °C with constant shaking at 200 rpm for 4 generations (selection time per experiment detailed in Table S1), allowing mutant library amplification and entry into exponential growth phase.

The competition cycle (output) was then initiated by inoculating cells from the input cycle into competition medium (SC-URA/MET/ADE supplemented with 200 μg mL^−1^ methotrexate) at a starting OD600 of 0.05. For this, a sufficient volume of cells was collected, centrifuged at 3,000 rpm for 5 minutes, and resuspended in pre-warmed output medium. Concurrently, each input replicate culture was split in half, harvested by centrifugation at 5,000 g for 5 minutes at 4 °C, washed with water, and the pellet stored at −20 °C for subsequent DNA extraction.

After approximately 4 generations in the competition cycle, each output replicate culture was similarly split in half, collected by centrifugation at 5,000 g for 5 minutes at 4 °C, washed twice with water, and the pellet stored at −20 °C.

#### DNA Extraction and Plasmid Quantification

The following DNA extraction protocol was optimized for a 100 mL culture at OD₆₀₀ ≈ 1.6. The protocol was scaled proportionally based on the harvested culture volumes for each screening experiment, as detailed in Table S1. Cell pellets from each input/output experimental replicate were resuspended in 1 mL of DNA extraction buffer. The suspensions were subjected to two freeze-thaw cycles, each involving freezing in a dry ice-ethanol bath followed by incubation in a 62 °C water bath. Subsequently, 1 mL of phenol/chloroform/isoamyl alcohol (25:24:1, pre-equilibrated with 10 mM Tris-HCl, 1 mM EDTA, pH 8.0) and 1 g of acid-washed glass beads (Biospec) were added. The samples were vigorously vortexed for 10 minutes and then centrifuged at 4,000 rpm for 30 minutes at room temperature. The supernatant was collected into a new tube, and this extraction step was repeated twice.

To the combined aqueous phase, 0.1 mL of 3 M sodium acetate (NaOAc) and 2.2 mL of ice-cold absolute ethanol were added. The mixture was gently inverted and incubated at −20 °C for 30 minutes. DNA was pelleted by centrifugation at maximum speed for 30 minutes at 4 °C. After discarding the ethanol supernatant, the DNA pellet was air-dried overnight at room temperature.

The dried DNA pellet was resuspended in 0.6 mL of 1× TE buffer, followed by the addition of 5 µL of RNase A (10 mg mL^−1^, Solarbio) and incubation at 37 °C for 30 minutes to digest RNA. The DNA solution was subsequently desalted and concentrated using the QIAEX II Gel Extraction Kit, with 50 µL of QIAEX II beads added. The beads were washed twice with PE buffer, and the DNA was finally eluted in 250 µL of 10 mM Tris-HCl (pH 8.5).

Plasmid concentration in the total DNA extracts, which contained yeast genomic DNA, was quantified by qPCR using primer pair oGJJ152/oGJJ153, which targets the plasmid origin of replication (ori) region.

#### Sequencing Library Construction

Sequencing libraries were constructed through two successive PCR amplifications. The first PCR (PCR1) amplified the target mutant proteins while incorporating frameshift nucleotides between the adapters and the sequencing region to enhance the diversity of initial sequencing bases. The second PCR (PCR2) introduced the remaining Illumina adapter sequences and sample-indexing barcodes.

To minimize PCR bias, the number of plasmid templates added to PCR1 was 20-50 times the expected number of sequencing reads. The plasmid template concentration did not exceed 6.25 × 10^7^ molecules μL^−1^ per reaction to avoid excessive co-amplification of yeast genomic DNA, which could interfere with PCR efficiency. PCR1 was performed using Q5 Hot Start High-Fidelity DNA Polymerase (New England Biolabs) according to the manufacturer’s instructions, with 25 pmol of a primer mixture (frameshift primers: oWCC144/103 for block 1; oWCC104/105 for block 2; oWCC106/145 for block 3). Forward and reverse primers were pooled based on oligonucleotide base diversity (see Table S1). Amplification was carried out for 10 cycles with an annealing temperature of 60 °C and an extension time of 20 seconds.

The PCR1 products were treated with ExoSAP-IT (Affymetrix) to remove residual primers, using 0.04 μL per μL of PCR reaction, incubated at 37 °C for 30 minutes, followed by enzyme inactivation at 80 °C for 15 minutes. Reactions for each sample were pooled and purified using the MinElute PCR Purification Kit (QIAGEN) according to the manufacturer’s protocol. DNA was eluted in EB buffer in a final volume equivalent to one-fifth of the total PCR1 volume.

PCR2 was performed separately for each sample using Hot Start High-Fidelity DNA Polymerase (NEB). The total PCR2 volume was one-quarter of the PCR1 volume, with PCR1 product added at a ratio of 0.05 μL per μL of reaction. This round incorporated the remaining portions of the Illumina adapters. All samples used the same forward primer (containing the 5’ P5 adapter), while the reverse primers (containing the 3’ P7 adapter) included unique barcode sequences (Table S1) to enable sample multiplexing (for indexes used per replicate in each sequencing run, see Table S1). PCR2 was performed for 12 cycles with an annealing temperature of 62 °C and an extension time of 20 seconds.

Reactions for each sample were pooled, and an aliquot was analyzed and quantified on a 2% agarose gel. Samples with different barcodes were mixed in equimolar ratios prior to sequencing. The pooled library was gel-purified and extracted using the QIAEX II Gel Extraction Kit (QIAGEN). The final constructed sequencing library was sequenced on an Illumina platform using 150 bp paired-end sequencing, performed by the CRG Genomics Core Facility and Baiteng Biotechnology.

#### Sequencing Data Processing

Paired-end FastQ files from all BindingPCA and AbundancePCA experiments were processed using DiMSum v1.2.9(*4*) (https://github.com/lehner-lab/DiMSum) with default settings and minor adjustments. Table S2 contains the DiMSum fitness estimates and associated errors for all experiments. The experimental design files and command-line options required to run DiMSum on these datasets are available on GitHub (https://github.com/weng-lab-ustc/Multimodal-allostery). In all cases, an adaptive minimum input read count threshold based on the number of corresponding nucleotide substitutions (the “fitnessMinInputCountAny” option) was selected to minimize the read fraction per variant associated with “variant flow” from sequencing error-induced low-order mutants.

Variant counts associated with all samples (output from DiMSum stage 4) were further filtered using custom scripts to retain variants consistent with the design of the respective libraries. For the nicking library (block 1), double-amino-acid variants were retained only if they contained one of the identified background mutations(*5*) : T2K, L6H, G12D, G12V, G12C, V14S, S17N, L19P, T35S, E37G, D38C, Y40A, Q61L, E63V, or Y64A. For the synthetic library (blocks 2 and 3), double-amino-acid variants were retained only if they contained the fixed background mutations specified by the library design, namely N116 and S145, respectively. Reads associated with double-amino-acid variants that did not contain the designated background mutation were discarded. Finally, fitness estimates and associated errors were derived from the resulting filtered variant counts using DiMSum (countPath option).

#### Thermodynamic Model Fitting with MoCHI

We used MoCHI v0.9 (https://github.com/lehner-lab/MoCHI) to fit global mechanistic models simultaneously to all 27 deep mutational scanning (ddPCA) datasets (9 phenotypes × 3 blocks). Briefly, individual KRAS protein-protein interactions (PPIs) were modeled as a three-state equilibrium—unfolded/unbound (*uu*), folded/unbound (*fu*), and folded/bound (*fb*)—assuming negligible unfolded/bound occupancy and additive residue-specific contributions to folding (Δ*G*_f_) and binding (Δ*G*_b_) free energies. Binding energies were interaction-specific, whereas folding energy was intrinsic to KRAS. Mutational effects on abundance were primarily attributed to folding changes, while other cellular factors were acknowledged as potential modifiers.

The MoCHI neural network comprises nine additive latent trait layers (eight binding traits and one folding trait), together with linear transformation layers that map latent energies to experimental readouts across assays (five AbundancePCA and 30 BindingPCA measurements). Nonlinear thermodynamic transformations (‘TwoStateFractionFolded’ and ‘ThreeStateFractionBound’) convert free energy estimates into folded and bound fractions, respectively.

The model was trained on a total of 35 ddPCA datasets, including 14 datasets generated in this study and 21 datasets previously reported by Weng *et al*. Training data included fitness measurements for wild-type, single, and double amino acid variants across all datasets.

During training, 30% of double-substitution variants were held out, with 20% for validation and 10% for testing. Validation loss guided hyperparameter selection, and early stopping was applied when the standard deviation of wild-type free energies over 10 epochs fell below 10^−3^. Model parameters θ were optimized via stochastic gradient descent using a weighted, regularized mean absolute error (MAE):

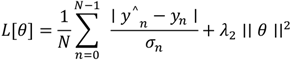

where *y_n_* and *σ_n_* are the observed fitness and its standard error, ŷ*_n_* is the predicted fitness, N is the batch size, and λ₂ = 10⁻^6^ penalized large free energy changes. Fitness errors weighted the MAE, and training data were resampled according to their error distributions; validation and test sets remained unchanged. Models were trained with the Adam optimizer (initial learning rate 0.025) for up to 1000 epochs, with exponential learning rate decay (γ = 0.98) if validation loss plateaued.

Free energies were computed from model parameters as Δ*G*_b_ = *θ_b_RT* and Δ*G*_f_ = *θ_f_RT* (*T* = 303 K, *R* = 0.001987 kcal K^−1^ mol^−1^). Confidence intervals were estimated via Monte Carlo simulations over ten independent model fits, using random train-validation-test splits and fitness resampling. Variants with 95% confidence intervals < 1 kcal mol^−1^ were considered robust. Inferred folding and binding free energies for all variants are provided in Table S3.

#### Data Normalization and Processing

Raw sequencing data from the three mutagenesis libraries (blocks) were initially used to calculate the growth rate and fitness for each mutant. To ensure comparability of results across different experimental libraries, the measured values for wild-type (WT) and stop codon (STOP) mutants were weighted and averaged, and normalization was performed using linear regression. Specifically, the growth rate and fitness data for each library were calibrated against the benchmark points of STOP and WT mutants, mapping them to a unified standardized scale to obtain normalized estimates (normalized growth-rate and normalized fitness), while simultaneously propagating the associated errors.

For fitness heatmap visualization, only single amino acid substitutions were retained. For replicate measurements of the same amino acid sequence appearing in different experiments, the values were consolidated using an error-inverse weighting method to derive consensus fitness and growth rate estimates for each unique mutant.

Finally, by aligning with the wild-type amino acid sequence, the specific location of each mutation on the KRAS protein was determined, and standard mutation nomenclature (e.g. G12D, Q61H) was generated. Through this series of processing steps, we obtained a standardized, non-redundant dataset of mutational effects with positional annotation, suitable for comparative analysis and structural mapping.

#### Enrichment analysis

Structural and functional enrichment of specific mutation classes or residue groups was evaluated using two-sided Fisher’s exact tests. For each analysis, residues or mutations were classified according to whether they belonged to the structural or functional category of interest, and enrichment was assessed using a 2 × 2 contingency table. Odds ratios (ORs) were calculated to quantify the magnitude of enrichment, with *p* values obtained from two-sided Fisher’s exact tests.

---

**Fig. S1.**
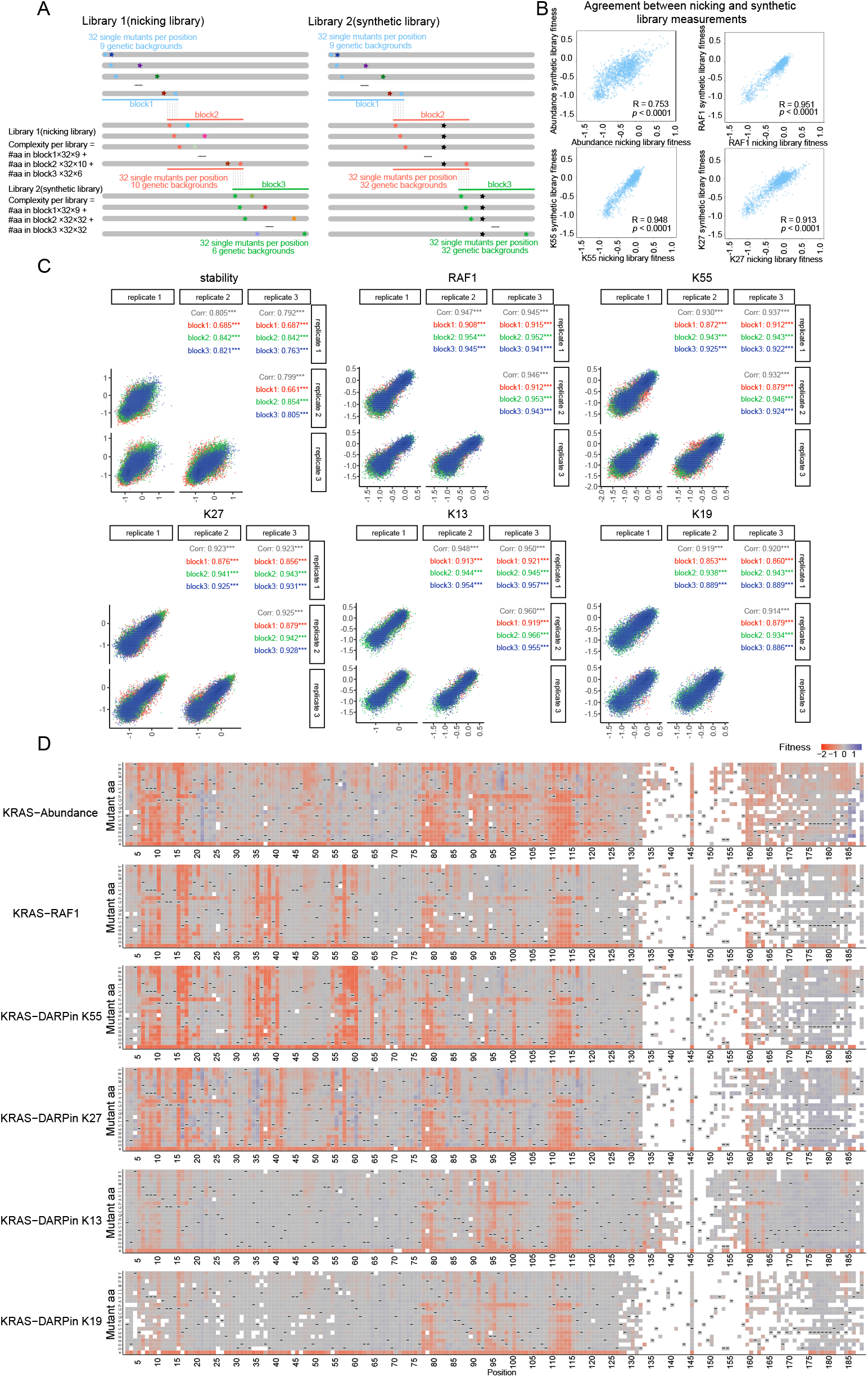
Library composition and sequencing data evaluation. (A) Composition of the nicking-based and synthetic KRAS mutagenesis libraries. Together, the two complementary libraries comprised over 80,000 KRAS variants, including more than 3,000 single amino acid substitutions and over 77,000 double amino acid substitutions (> 49,000 from the synthetic library and > 28,000 from the nicking library). In this study, the nicking-mutagenesis library covered block 1 of KRAS, whereas the synthetic library covered blocks 2 and 3. Binding to RAF1-RBD, DARPins K55, K27, K13, and K19, together with folded protein abundance, was quantified in triplicate using the synthetic library. Binding of block 1 variants to DARPins K13 and K19 was measured using the nicking-mutagenesis library. (B) Comparison of abundance and RAF1/K55/K27 binding fitness measurements obtained from the nicking and synthetic libraries for single amino acid substitutions across KRAS block 2 and block 3. (C) Pearson correlations between biological replicates for KRAS abundance and binding fitness measurements with RAF1, K55, K27, K13, and K19 in nicking library (block 1) and synthetic library (block 2 and block 3). (D) Heatmaps showing fitness effects of single amino acid substitutions on KRAS abundance and binding to RAF1, K55, K27, K13, and K19, as measured by AbundancePCA (top) and BindingPCA (bottom). Empty cells indicate missing data; dashes denote wild-type residues; asterisks indicate stop codons. To complete mutational coverage across KRAS, these measurements were integrated with our previously published block 1 dataset for RAF1-RBD, K55, K27, and abundance(*5*).

**Fig. S2.**
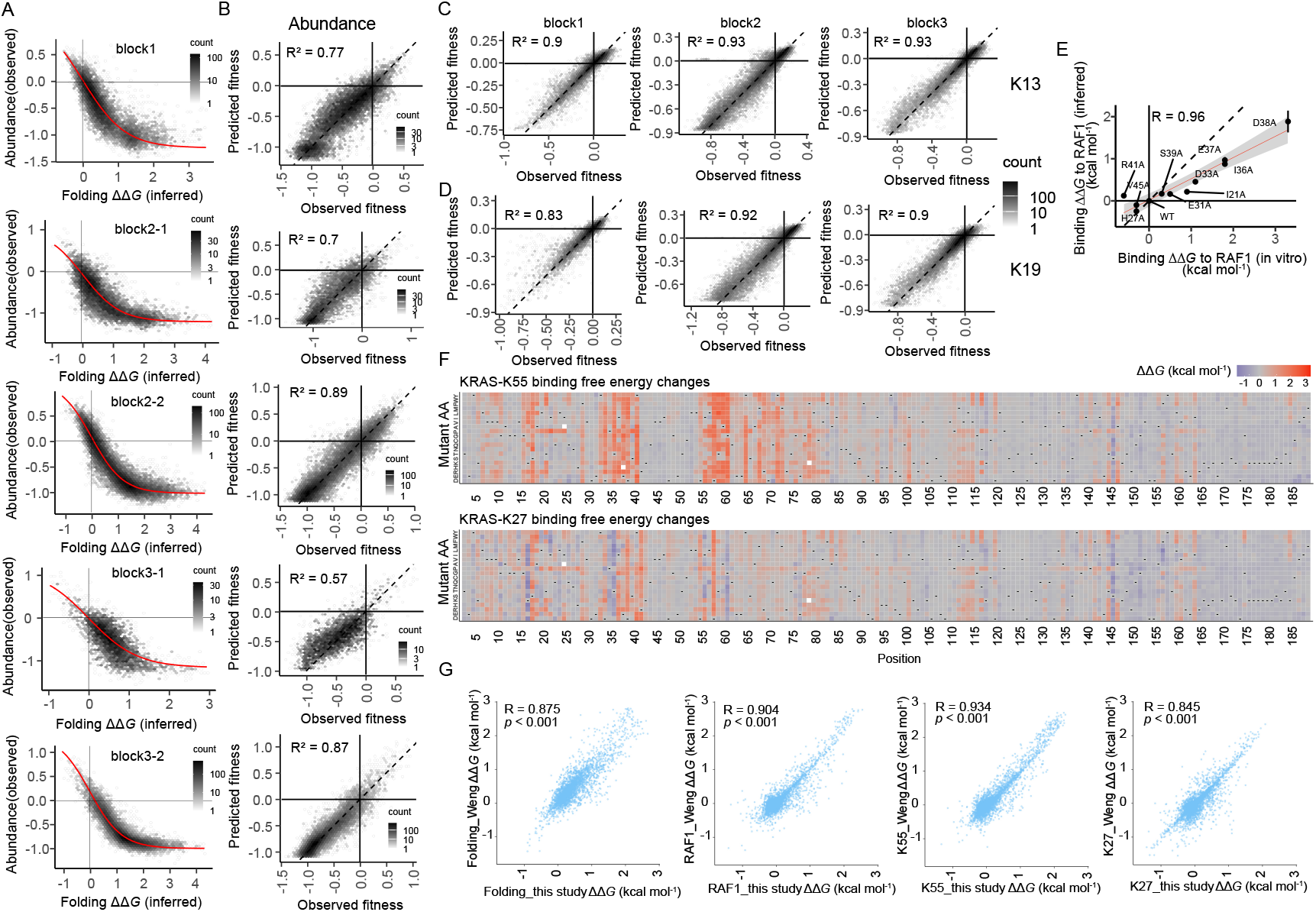
MoCHI model evaluation and energy heatmaps. (A) Two-dimensional density plot illustrating the non-linear relationship (global epistasis) between observed AbundancePCA fitness and inferred folding free energy changes. (B) Two-dimensional density plot comparing MoCHI model predictions and observed AbundancePCA fitness for held-out test data (10% of double-amino acid substitution variants excluded from training). (C) Two-dimensional density plot comparing MoCHI model predictions and observed BindingPCA fitness for DARPin K13 in the held-out test data. (D) Two-dimensional density plot comparing Mochi model predictions and observed BindingPCA fitness for DARPin K19 in the held-out test data. (E) Comparisons of model-inferred free energy changes to *in vitro* measurements(*6*). Error bars indicate 95% confidence intervals from a Monte Carlo simulation approach (n = 10 experiments). Linear regression fit and its 95% confidence interval are shown as a red solid line and a grey shaded area, respectively. Pearson’s R is shown. Black dashed line indicates *y* = *x*. (F) Heatmaps showing inferred binding free energy changes for KRAS interactions with DARPins K55 and K27. (G) Correlation between binding free-energy changes inferred in this study and those reported for KRAS folding and binding to RAF1, K55, and K27 in Weng *et al*(*5*). In total, we quantified 26,778 binding free energy changes (ΔΔ*G*_b_) across eight binders (RAF1-RBD, RALGDS-RBD, PIK3CG-RBD, SOS1, and DARPins K55, K27, K13 and K19), of which 18,956 substitutions overlap with measurements reported by Weng *et al*. and 7,822 represent newly generated data. We also quantified 3,551 folding free energy changes (ΔΔ*G*_f_), of which 3453 substitutions overlap with the previous dataset and 98 are newly generated.

**Fig. S3.**
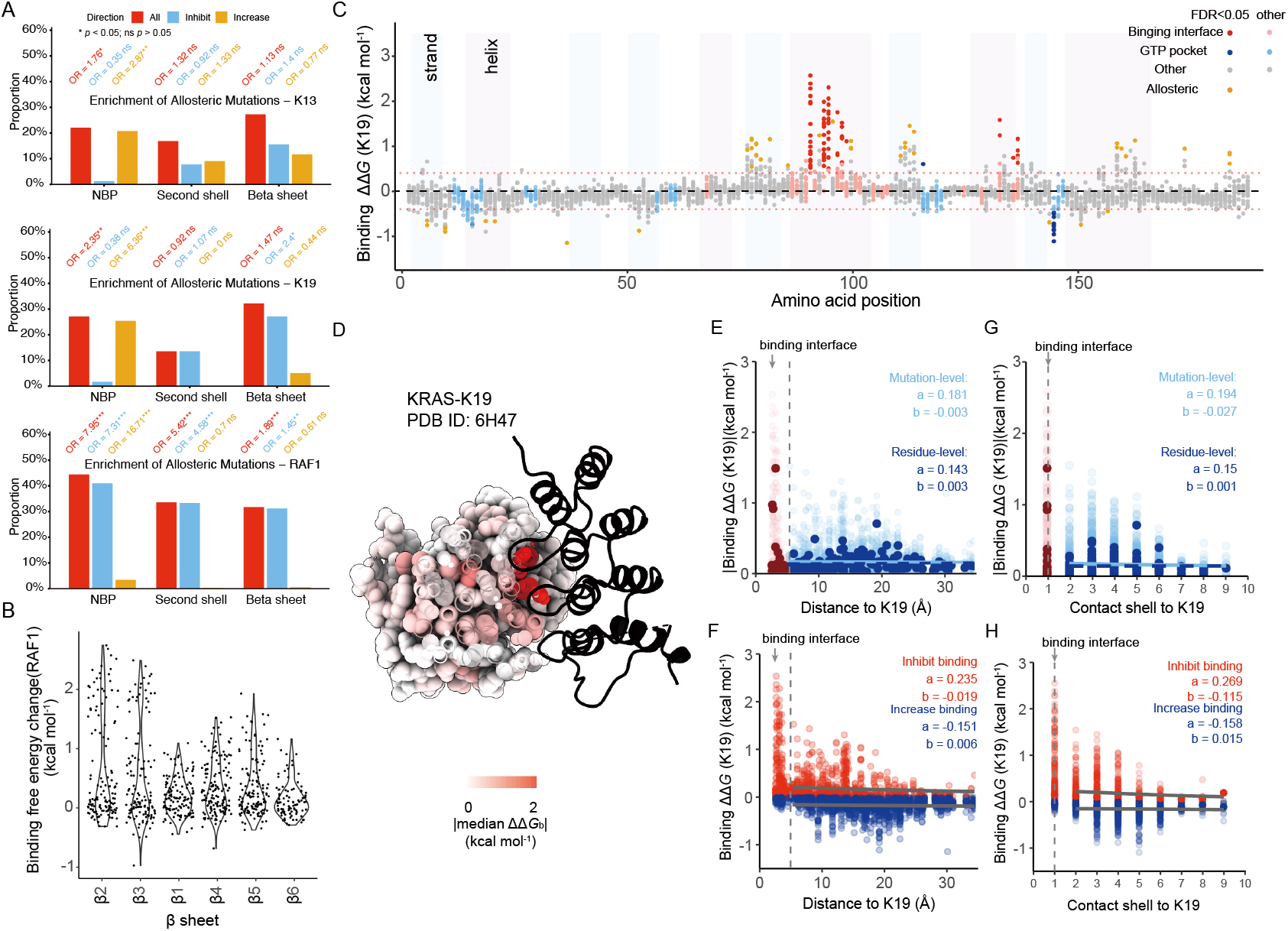
Comparative allosteric landscapes of two KRAS interfaces. (A) Enrichment analysis of allosteric mutations of K13, K19, and RAF1 in NBP, second shell, and β-sheet. (B) Violin plot showing the decay of binding free energy change across successive strands in the β-sheet. β-strands are ordered by increasing distance to RAF1 in the 3D structure. (C) Manhattan plot of binding free energy changes for all single amino acid substitutions. Points are colored by residue position; significance was assessed using a two-sided z-test with FDR = 0.05. (D) Structural visualization of the decay of |ΔΔ*G*_b_| (kcal mol^−1^) with increasing distance from the ligand. KRAS bound to the indicated ligand is shown, colored by median absolute ΔΔ*G*_b_ (kcal mol^−1^) values. (E) Relationship between absolute ΔΔ*G*_b_ (kcal mol^−1^) of K19 and the minimum distance from side-chain heavy atoms to the ligands. Exponential decay fits (*y* = *a* · *e^bx^*) are applied to both mutation-level |ΔΔ*G*_b_| (kcal mol^−1^) values and residue-level median |ΔΔ*G*_b_| (kcal mol^−1^) values. (F) Relationship between ΔΔ*G*_b_ (kcal mol^−1^) of K19 mutations and the minimum distance from side-chain heavy atoms to the ligand. Mutations are stratified into inhibit (ΔΔ*G*_b_ > 0) and increase (ΔΔ*G*_b_ < 0) groups. Exponential decay models (*y* = *a* · *e^bx^*) are independently fitted to mutation-level ΔΔ*G*_b_ values for each group. (G) Relationship between absolute ΔΔ*G*_b_ (kcal mol^−1^) of K19 and the contact shell to the ligands. Exponential decay fits (*y* = *a* · *e^bx^*) are applied to both mutation-level ΔΔ*G*_b_ (kcal mol^−1^) values and residue-level median ΔΔ*G*_b_ (kcal mol^−1^) values. (H) Relationship between ΔΔ*G*_b_ (kcal mol^−1^) of K19 and the contact shell to the ligands. Mutations are stratified into inhibit (ΔΔ*G*_b_ > 0) and increase (ΔΔ*G*_b_ < 0) groups. Exponential decay models (*y* = *a* · *e^bx^*) are independently fitted to mutation-level ΔΔ*G*_b_ values for each group.

**Fig. S4.**
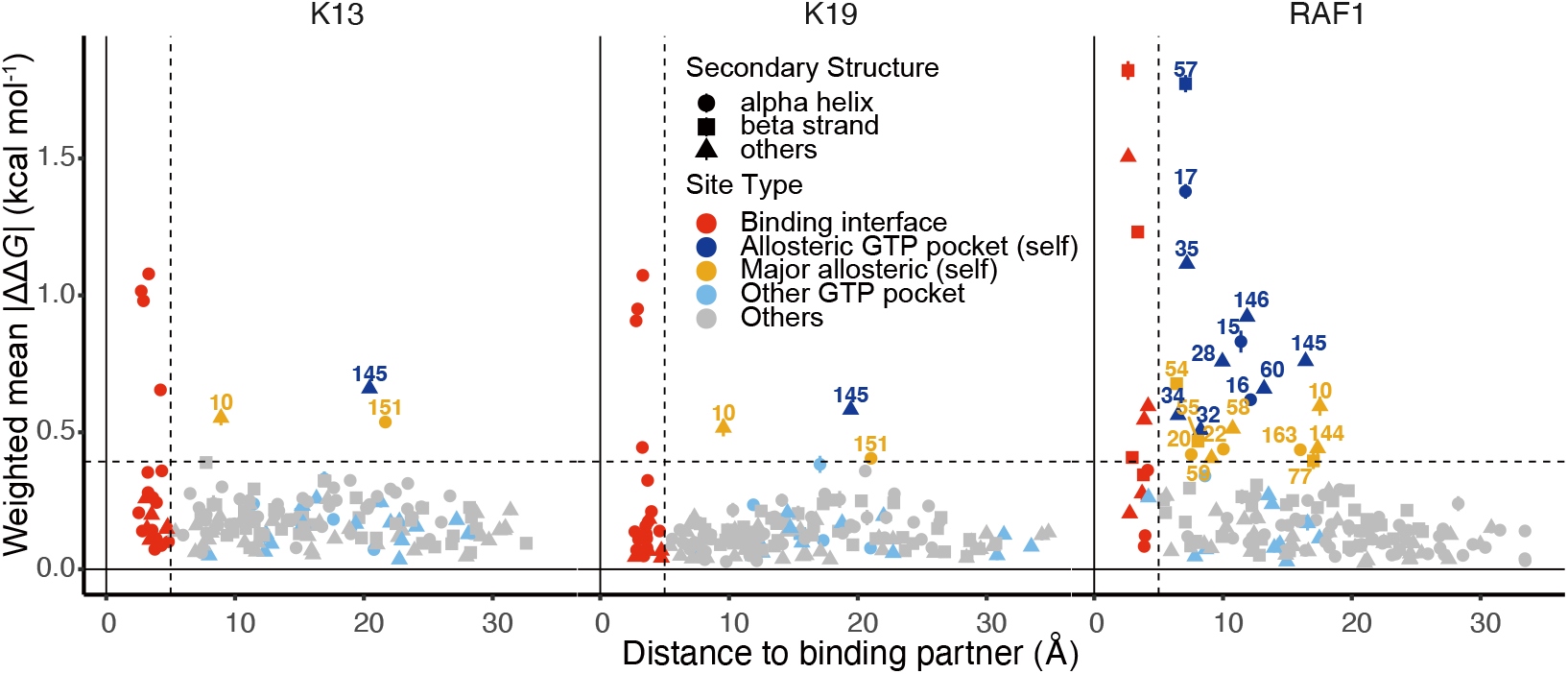
Comparative allosteric regulation of KRAS binding through two interfaces. The relationship between the site-averaged absolute change in binding free energy and the minimum distance from side-chain heavy atoms to their respective ligands for K13, K19, and RAF1. Major allosteric sites are defined as residues outside the binding interface whose weighted mean absolute change in binding free energy exceeds the average weighted mean absolute change in binding free energy of mutations at binding interface residues across the eight binders (indicated by the horizontal dashed line). Error bars represent 95% confidence intervals (n ≥ 10).

**Fig. S5.**
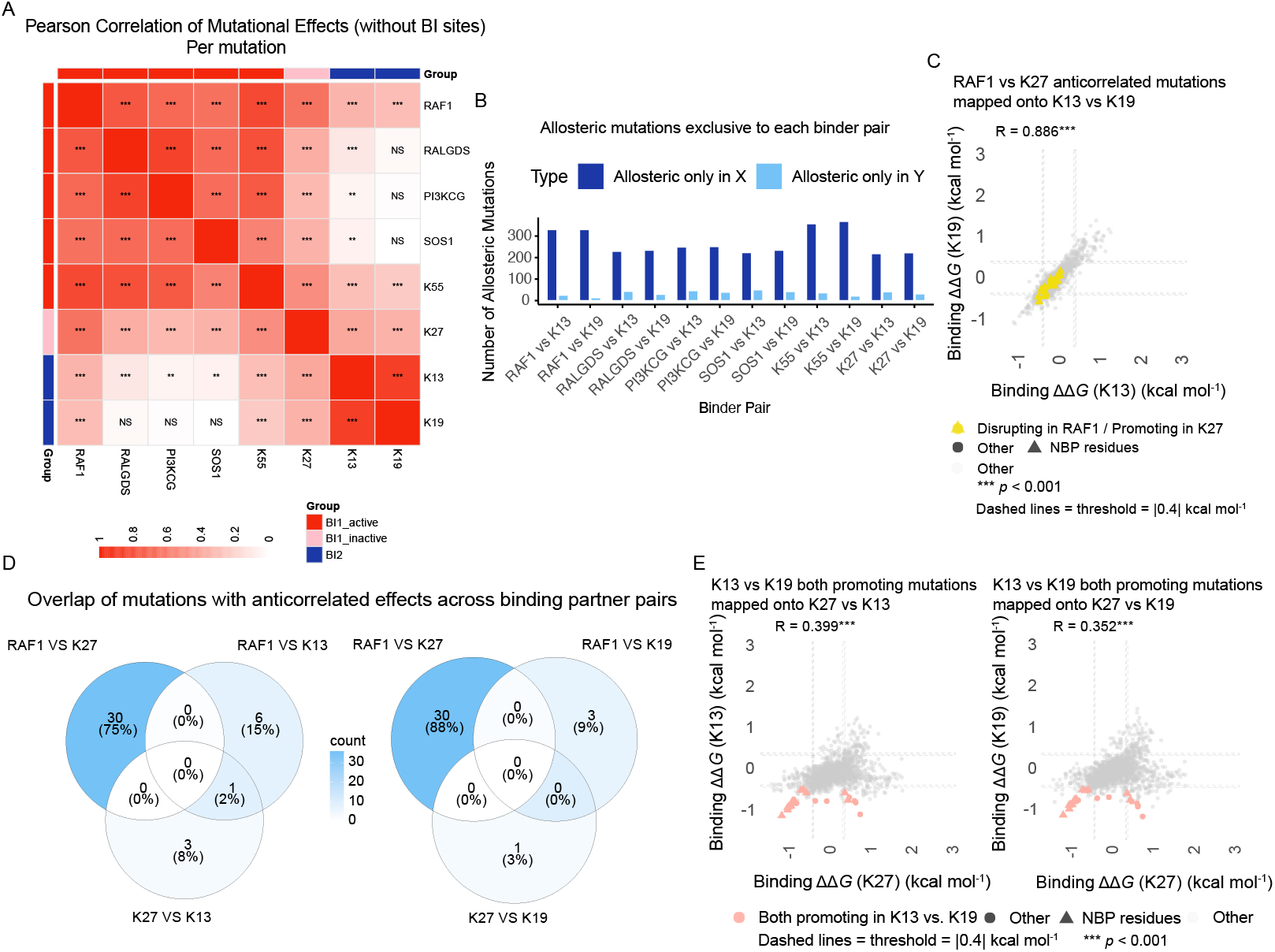
Three forms of allosteric regulation in KRAS. (A) Pearson correlation heatmap of mutational effects for eight binders pairwise comparison, excluding BI site mutations. Color gradient: white (correlation = 0) to red (positive correlation). Cell labels: *** (*p* < 0.001), ** (*p* < 0.01), * (*p* < 0.05), ns (not significant, *p* ≥ 0.05); diagonal blank. Row/column annotations: BI1 active binder (red), BI1 inactive binder (pink), BI2 binder (blue), BI1 denotes interface 1 binding, with “active” and “inactive” referring to the corresponding KRAS binding states; BI2 denotes interface 2 binding. (B) Bar plot displays the count of allosteric mutations uniquely identified in each binder of pairwise comparisons between BI1 binders (RAF1, RALGDS, PI3KCG, SOS1, K55, K27) and BI2 binders (K13, K19). Dark blue bars: allosteric mutations only in BI1 binder. Light blue bars: allosteric mutations only in BI2 binder. X-axis shows binder pairs ordered as indicated. Y-axis shows number of allosteric mutations. (C) Scatter plot of pairwise mutational effects between K13 and K19 (without binding interface sites). Each point represents a mutation; Mutations previously defined as KRAS state-switch (allosteric mutations with opposing effects on RAF1 and K27, promoting the binding of K27 but disrupting the binding of RAF1) are highlighted. Dashed lines indicate thresholds for effect directionality. (D) Venn diagram showing overlap among mutations exhibiting anticorrelated effects in three pairwise comparisons: RAF1 vs. K13/K19, RAF1 vs. K27, and K27 vs. K13/K19. Numbers indicate mutation counts in each intersection. (E) Scatter plots of pairwise mutational effects between K27 vs. K13/K19 (without binding interface sites). Each point represents a mutation. Highlighted mutations correspond to allosteric mutations that increase binding to both K13 and K19. Dashed lines indicate thresholds for effect directionality.

**Fig. S6.**
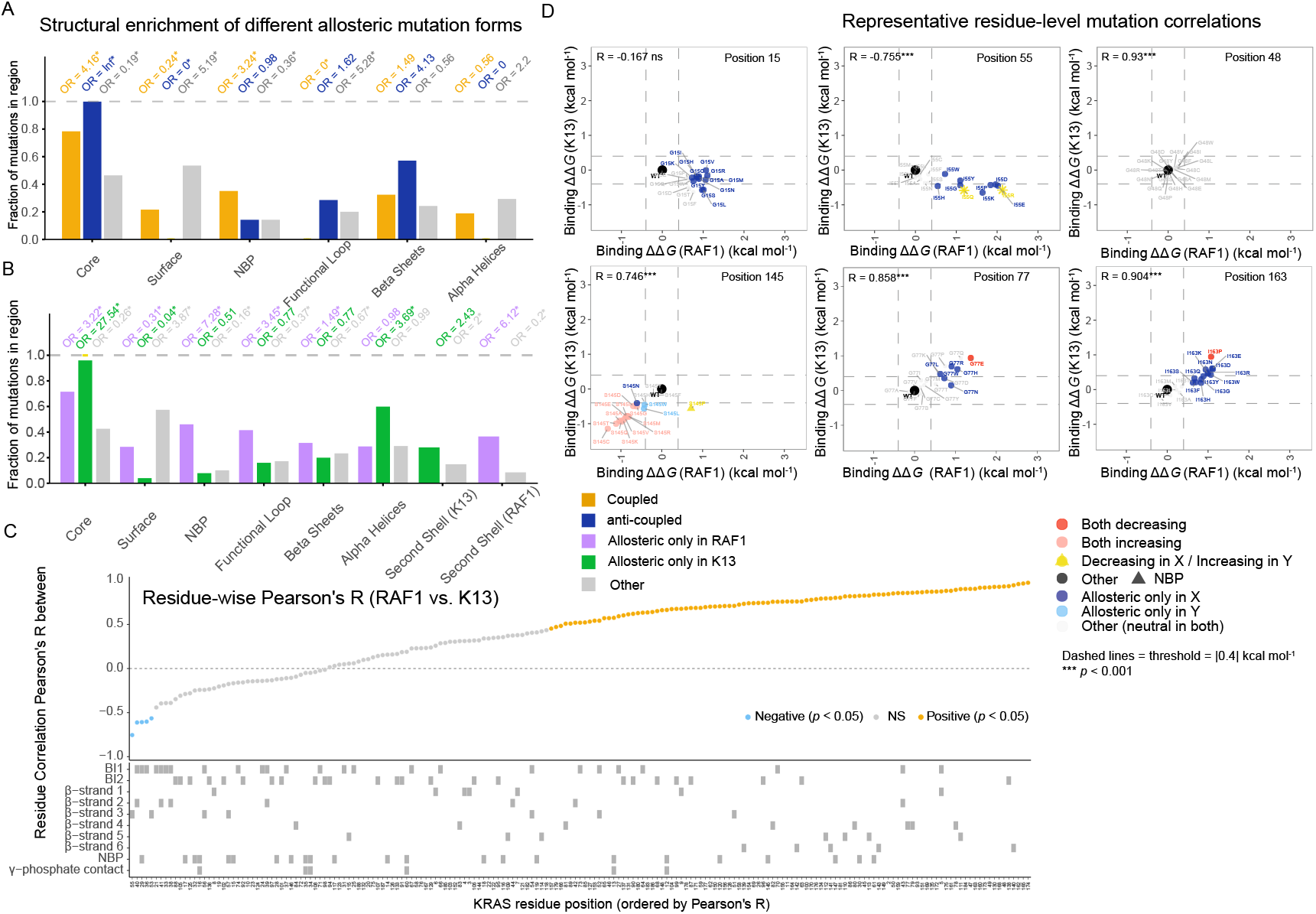
Coupled, anti-coupled and independent allostery across the KRAS structure. (A) Bar plots showing the fraction of allosteric mutations (coupled, anti-coupled, and other) located in core, surface, nucleotide-binding pocket (NBP), functional loop regions (switch I, switch II, and P-loop), β-sheet, and α-helix residues for three pairwise comparisons RAF1 vs. K13. Enrichment quantified using odds ratios (ORs) and Fisher’s exact test relative to all other mutations. Significant enrichment (*p* < 0.05) indicated by asterisks. (B) Bar plots show the fraction of RAF1 vs. K13 allosteric mutations (RAF1-specific, K13-specific, and other) located in core, surface, nucleotide-binding pocket (NBP), functional loop regions (switch I, switch II, and P-loop), β-sheet, α-helix and second-shell residues. Enrichment was quantified using odds ratios (ORs) and Fisher’s exact test relative to all other mutations. (C) Correlation coefficients (Pearson’s R) between ΔΔ*G*_b_ (RAF1) and ΔΔ*G*_b_ (K13) across 19 mutations per residue. Each data point represents one KRAS residue. Orange point: residues with positive correlation (*p* < 0.05). Light blue point: residues with negative correlation (*p* < 0.05); grey points indicate non-significant correlation (*p* > 0.05). (D) Scatter plot of ΔΔ*G*_b_ (RAF1) versus ΔΔ*G*_b_ (K13) for selected residues from Fig. S6C, showing 19 mutations and wild-type per residue. Each panel represents one selected residue. Each point represents a single mutation, x-axis: ΔΔ*G*_b_ (RAF1) (kcal mol^−1^); y-axis: ΔΔ*G*_b_ (K13) (kcal mol^−1^). All 19 mutations per residue are labeled individually. Point colors follow the allosteric mutation definition from Fig. 5A.

**Fig. S7.**
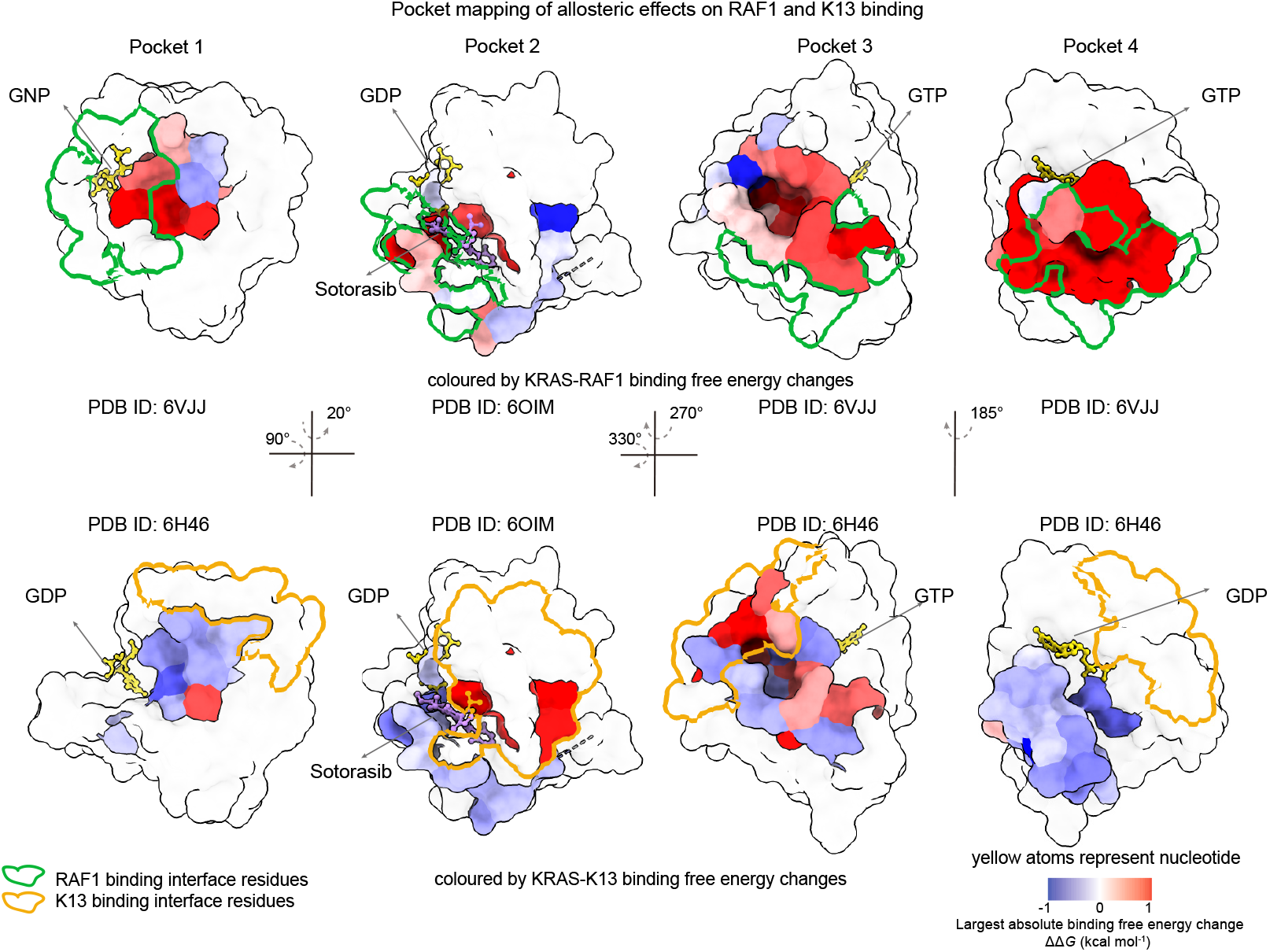
Multimodal allosteric regulation from the KRAS surface and pockets. Structural alignment of KRAS bound to RAF1 and K13 (PDB IDs: 6VJJ and 6H46, pocket 2 PDB IDs: 6OIM), with KRAS surface coloured using the previously described pocket annotation scheme. Residues are coloured according to the signed ΔΔ*G* values of mutations with the largest absolute binding free energy changes for RAF1 and K13 at each residue, the colour scale is capped at −1 and +1 kcal mol^−1^; values beyond this range are shown at the corresponding colour-scale limits. Pocket 1, residues 5–7, 39, 54–56 and 70–75; pocket 2, residues 10, 13, 17, 35, 59, 60–62, 64, 69, 70, 73, 93, 96, 97, 100, 101, 104 (Residues in contact with sotorasib in PDB ID 6OIM); pocket 3, residues 97, 101, 107–111, 136–140 and 161–166; pocket 4, residues 17, 21, 24–40 and 57. Green outlined surface represents interface 1; Orange outlined surface represents interface 2; arrows indicate the direction of rotation. The limits of the colour scale are defined for visualization and do not represent the maximum or minimum binding free energy change at each residue. The mutations are located in the HVR, which is absent from the crystal structure used for visualization and therefore are not displayed.

**Table S1.**

Experimental procedures, DiMSum analysis and processing, and primer sequences for the nicking library, plasmids, PCR1 and PCR2.

**Table S2.**

Fitness estimates for KRAS abundance and binding to six binding partners.

**Table S3.**

Inferred folding and binding free energy changes for KRAS variants.

**Table S4.**

Contacts at the interaction interfaces of DARPin K13 and K19 with KRAS.

**Table S5.**

Allosteric forms associated with mutations located on the surface and in the pockets of KRAS, in the context of RAF1 vs. K13 results.

**Movie S1. 3D structures of KRAS in complex with RAF1, DARPin K13 and DARPin K19.**

3D structures of KRAS in complex with RAF1, DARPin K13 and DARPin K19 (PDB IDs: 6VJJ, 6H46 and 6H47). (Video corresponding to Fig. 1D.)

**Movie S2. 3D structure of KRAS bound to K13 and K19.**

3D structure of KRAS bound to DARPin K13 and K19 (PDB ID: 6H46 and 6H47). Residues are coloured according to the position-wise median ΔΔ*G*_b_ (kcal mol^−1^). DARPin K13 and K19 are shown as grey ribbons. (Video corresponding to Fig. 2B)

**Movie S3. 3D structure of KRAS bound to K13, K19, and RAF1.**

3D structure of KRAS bound to DARPin K13, K19, and RAF1 (PDB ID: 6H46, 6H47 and 6VJJ) in which residue atoms are coloured according to the position-wise median ΔΔ*G*_b_ (kcal mol^−1^) for binding. (Video corresponding to Fig. 3B and Fig. S3D.)

**Movie S4. 3D structure of KRAS bound to K13, K19, and RAF1.**

3D structure of KRAS bound to DARPin K13, K19, and RAF1 (PDB IDs: 6H46, 6H47 and 6VJJ) in which residues are coloured according to the position-wise weighted mean ΔΔ*G*_b_ (kcal mol^−1^) for binding. Green outlines indicate residues identified as allosteric hotspots in each binder. (Video corresponding to Fig. 4A)

**Movie S5. 3D structures of KRAS bound to RAF1 or K13 showing allosteric mutation categories.**

3D structures of KRAS bound to RAF1 (PDB ID: 6VJJ) or DARPin K13 (PDB ID: 6H46) showing coupled, inversely(anti)-coupled, or independent mutations, corresponding to the six panels in Fig. 6A (left, middle, and right columns; top, RAF1; bottom, K13). Mutations are colored according to the position-wise ΔΔ*G*_b_ (kcal mol^−1^) or weighted mean ΔΔ*G*_b_ (kcal mol^−1^) for the corresponding binder. Only residues containing at least one mutation classified into the corresponding allosteric category are colored. When multiple mutations are present at the same residue, weighted mean ΔΔ*G*_b_ values are used. (Video corresponding to Fig. 6A)

**Movie S6. 3D structure of KRAS (Apo).**

3D structure of KRAS (PDB ID: 4OBE) showing the groups of residues defined in Fig. S6C. Orange, positive correlation residues supporting allostery (*p* < 0.05); green, negative correlation residues supporting allostery (*p* < 0.05); light grey, other correlation residues supporting allostery (*p* > 0.05); black, residues in the union of BI1 (RAF1/K55/K27; top) or BI2 (K13/K19; bottom); white, residues without allostery (residues supporting allostery indicate that the residue has a significant allosteric mutation (defined in Fig. 5A). (Video corresponding to Fig. 6C)

**Movie S7. 3D structure of KRAS (Apo).**

3D structure of KRAS (PDB ID: 4OBE) in which residues are coloured according to its correlation coefficient (Pearson’s R value) in Fig. S6C. The color scale indicates the range of R values. (Video corresponding to Fig. 6D)

